# Common and Sex-Biased Muscle Transcriptomic Aging Signatures Across Diverse Species

**DOI:** 10.64898/2026.10.16.759636

**Authors:** Eric C. Randolph, Yifei Fang, Cailin Kessen, Richard P. Meisel, James Walters, Tony Gamble, Gerald Wilkinson, Erica Larschan, Ashley Webb, IISAGE Consortium, Peggy R Biga, Nicole C. Riddle, Jingyue E. Duan

## Abstract

Sex differences in lifespan are widespread yet inconsistent across the animal kingdom. In mammals, XX females outlive XY males in over 60% of species, while this pattern reverses in birds, where ZZ males have a higher life expectancy. These sex differences raise the question of whether sex-biased aging biomarkers (age-associated molecular signatures that reflect biological aging processes) can be identified based on transcriptomic data and whether such markers are shared across taxa. To identify sex-biased aging biomarkers, we analyzed sex-specific skeletal muscle transcriptomes (RNA-seq) from young and old individuals of six species spanning major evolutionary lineages. Four female-biased and 18 male-biased aging biomarkers were identified across all species. Three shared aging biomarkers were also identified in both sexes of all species: Na Channel, Heat Shock Protein 70 (HSP70), and Secreted Protein Acidic and Rich in Cysteine (SPARC). Particularly, age-associated changes in the HSP70 family are suggestive of a shared decline in proteostasis, a hallmark of aging observed across both model and non-model organisms. Furthermore, using an Ornstein-Uhlenbeck (OU) model applied to orthogroup phylogenies, we identified lineage- and sex-dependent expression shifts in shared aging biomarkers. In addition, sex-biased orthogroups revealed age-associated alterations in muscle structure and function in females, whereas mitochondrial and metabolic pathways were predominantly changed in males, indicating divergent mechanisms in muscle aging between sexes. Together, these findings identified a set of transcriptional aging signatures, both shared and sex-biased, that are detectable across diverse evolutionary lineages, providing insight into shared and divergent molecular mechanisms underlying aging.

**Significance statement:** Across the animal kingdom, there are many instances of sex differences in longevity, suggesting the presence of sex differences in the underlying biological processes that drive longevity. This study investigated gene expression differences occurring in age among six species spanning large evolutionary time scales to investigate whether sex differences in biological processes are shared across evolution and whether they can serve as identifiers of aging biomarkers. This investigation yielded three ubiquitously shared transcriptional aging biomarkers along with four female-biased and 18 male-biased transcriptional aging biomarkers. These findings reveal a semi-concerted evolutionary aging pattern that aligns well with preestablished aging biomarkers. In addition, these findings reveal a sex-biased evolutionary aging pattern.

## Introduction

Across the animal kingdom, males and females frequently differ in lifespan (Bronikowski et al 2022), and these patterns vary systematically across major clades. In mammals, females outlive males in roughly 60% of surveyed species (Lemaître et al 2020), whereas in birds the opposite pattern is common, with males typically living longer (Marais et al 2018). These differences between birds and mammals often are interpreted in light of the heterogametic sex hypothesis, which posits that the sex possessing two different sex chromosomes (ZW females in birds, XY males in mammals) experiences greater deleterious effects associated with hemizygosity (Marais et al 2018, Staerk et at 2025). Fish also exhibit sex-biased longevity, although the direction of this bias varies by species. For example, male yellow tang (*Zebrasoma flavescens*) outlive females (Claisse et al 2009), while the opposite pattern is observed in the Maori snapper (*Lutjanus rivulatus*) (Wakefield et al 2020) and African annual turquoise killifish (*Nothobranchius furzeri*) (Reichard et al 2021). Many invertebrates experience sex-biased longevity as well, including *Drosophila melanogaster* (Iliadi et al 2009, Millington and Rideout 2018) and *Caenorhabditis elegans* (McCulloch and Gems 2003, Barr et al 2018). These widespread yet inconsistent patterns strongly suggest that intrinsic, sex-specific mechanisms are influencing aging; however, the underlying molecular basis of these differences remains poorly understood.

While substantial progress has been made in defining common biological processes underlying aging, further work is needed to determine the extent to which these mechanisms are shared across species and between sexes. The updated *Hallmarks of Aging* framework (López-Otín et al 2023) outlines key processes, such as loss of proteostasis, mitochondrial dysfunction, and genomic instability, that are thought to contribute broadly to organismal aging. However, these hallmarks were largely derived from studies in a limited set of classical model organisms, including *D. melanogaster*, *C. elegans*, and *Mus musculus* (Biga et al 2025). Importantly, many of the studies informing this framework either excluded one sex or were conducted predominantly in males, introducing a bias that may obscure sex-specific aging mechanisms (Biga et al 2025). This limitation reflects a broader and well-documented trend in biomedical research, where sex bias historically has been prevalent (Beery and Zucker 2011, Holdcroft 2007, Ballantyne and Rogers 2011) and continues to persist despite increased awareness (Carmody et al 2022). Consequently, it remains uncertain whether the established hallmarks of aging adequately reflect the full molecular diversity of aging across sexes and species, as current knowledge is disproportionately derived from model organisms and male-biased studies.

Comparative transcriptomics provides a powerful framework for determining whether the molecular processes underlying aging, and particularly sex-specific aging, are shared evolutionarily. By incorporating species that span deep phylogenetic divergence, this approach enables the identification of transcriptional signatures that are either ubiquitous across taxa or restricted to specific lineages or sexes. Analyzing these data with an Ornstein-Uhlenbeck (OU) modeling framework further enables the detection of adaptive expression shifts and evolutionary divergence along a phylogeny (Bedford and Hartl 2009). Despite these advantages, most prior transcriptomic studies of aging have focused on within-species analyses. Such studies, including large-scale fly aging atlases (Lu et al 2023), have been highly effective at identifying age-and sex-associated transcriptional changes within individual organisms. However, they provide limited insight into whether these patterns are shared across species or represent lineage-specific changes. As a result, there remains a critical gap in aging biology: the lack of rigorous, cross-species comparative analyses that systematically evaluate shared versus divergent transcriptomic hallmarks of aging across both model and non-model organisms.

Muscle (skeletal muscle in vertebrates and striated muscle in invertebrates) represents a particularly informative tissue for interspecific comparisons of aging. It is highly conserved in both structure and function across metazoans (Augustin and Partridge 2009, Mukund and Subramaniam 2019, Mendoza et al 2023), undergoes well characterized age-related decline (Tieland et al 2018), and is sensitive to systemic and sex hormone–mediated aging processes (Sipilȁ et al 2013, Velders and Diel 2013, Critchlow et al 2025). Importantly, muscle can be sampled in a minimally invasive manner and allows broad inclusion of both model and non-model species, including taxa for which other tissues cannot be ethically or feasibly obtained.

Here, we perform a comparative, cross-species transcriptomic analysis that integrates both model and non-model organisms while explicitly incorporating both sexes to assess whether aging signatures exhibit sex bias and whether they are species-specific or shared across evolutionary lineages. Using muscle RNA-seq datasets from six species, we first characterize sex-associated transcriptional changes within species during aging. We then evaluate the extent to which molecular pathways show shared age-related regulation across taxa and applied an OU modeling framework to identify lineage- and sex-biased expression shifts in key aging-associated gene families. Our analyses reveal a combination of shared, lineage-specific, and sex-biased transcriptional signatures of aging, identifying a set of aging biomarkers that are detectable across diverse evolutionary lineages. These findings provide insight into both shared and divergent molecular mechanisms underlying sex-specific aging.

## Results

### Sex and Age are Distinct Drivers of Expression Variation

To determine whether sex-biased aging biomarkers can be identified across a broad phylogenetic spectrum, and whether such markers are shared across major animal clades, we integrated previously published RNA-seq datasets with newly collected datasets. These datasets included young (sexually mature) and old (post-median lifespan) individuals from six species spanning deep evolutionary divergence (**Fig. 1A-B**): two insects (*D. melanogaster* and *Plodia interpunctella)*, one fish (*N. furzeri*), and three mammals, including two rodents (*M. musculus* and *Rattus norvegicus*) and humans (*Homo sapiens*). Overall, these species span hundreds of millions of years of evolution while also including closely related pairs, enabling comparisons of aging signatures across both short and long evolutionary time.

**Figure 1.**
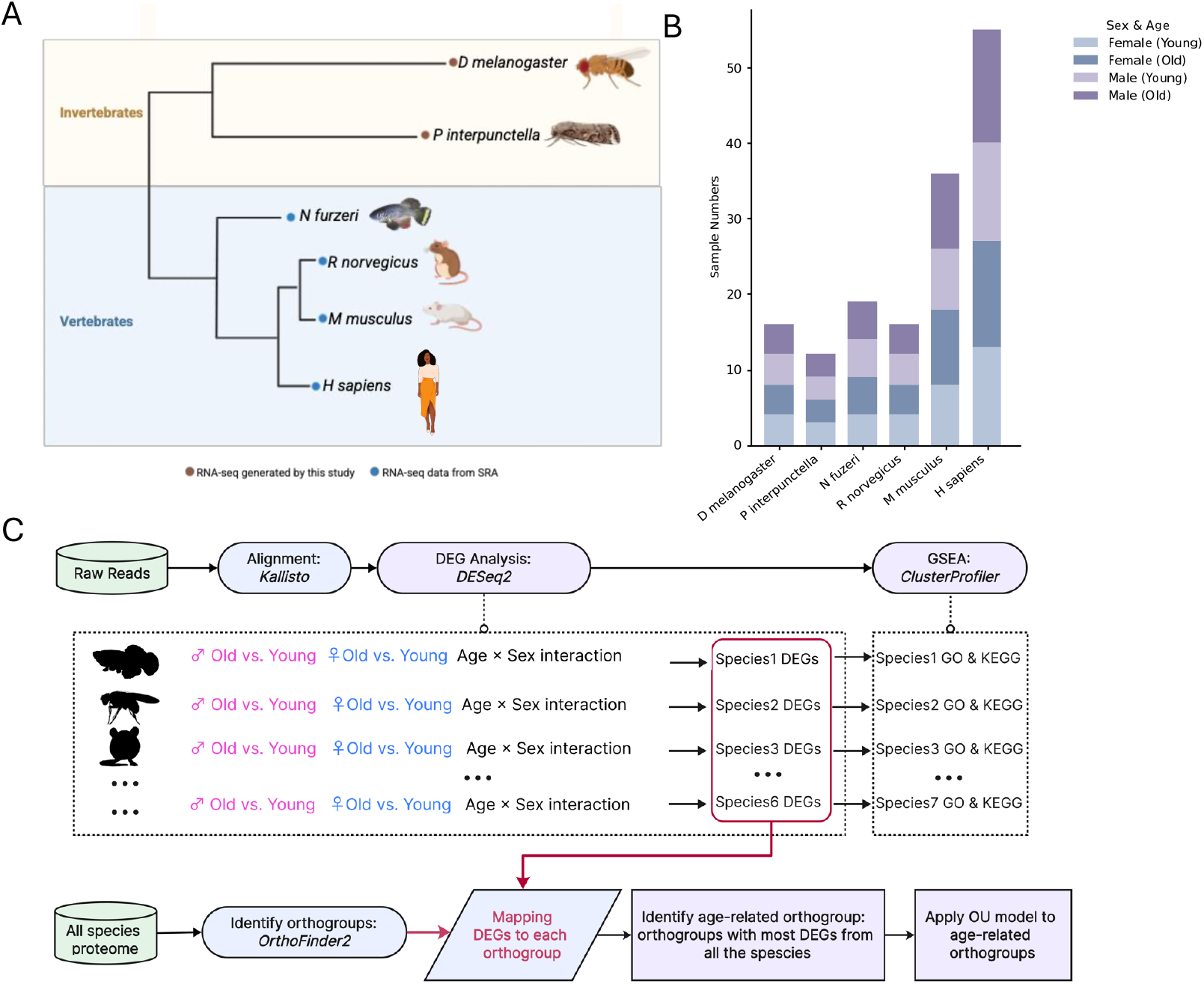
Overview of samples and study design. A) phylogenetic tree showing the evolutionary relationships of the species used in this study. The color of circles next to the species names are colored to indicate whether RNA-sequencing raw reads were generated by collaborators of this study (brown) or from previously published data retrieved from the Gene Expression Omnibus (GSO; blue). B) The total number of samples used for each species. Within each species, bars are colored according to the sex and age of that sample. C) Graphical illustration of the bioinformatic workflow which was designed for and used in this study. **Alt text:** Figures showing the evolutionary tree of the species studied in this project, the sample distribution for each species, and an overview workflow.

Following a standardized bioinformatics analysis pipeline (**Fig. 1C**), we performed principal component analysis (PCA) on normalized transcript counts within each species to assess whether transcriptional profiles could distinguish samples by age and sex. Distinct clustering by age and/or sex was observed in all species (**Fig. 2A**), indicating robust biological signals. Specifically, *P. interpunctella*, *R. norvegicus*, and *N. furzeri* showed a strong separation by age, whereas *D. melanogaster*, *R. norvegicus, M. musculus*, and *H. sapiens* exhibited a strong separation by sex. These results demonstrate that the dominant source of transcriptional variation is not universally driven by age; interestingly, in several species, sex was the primary determinant of transcriptomic differences.

**Figure 2.**
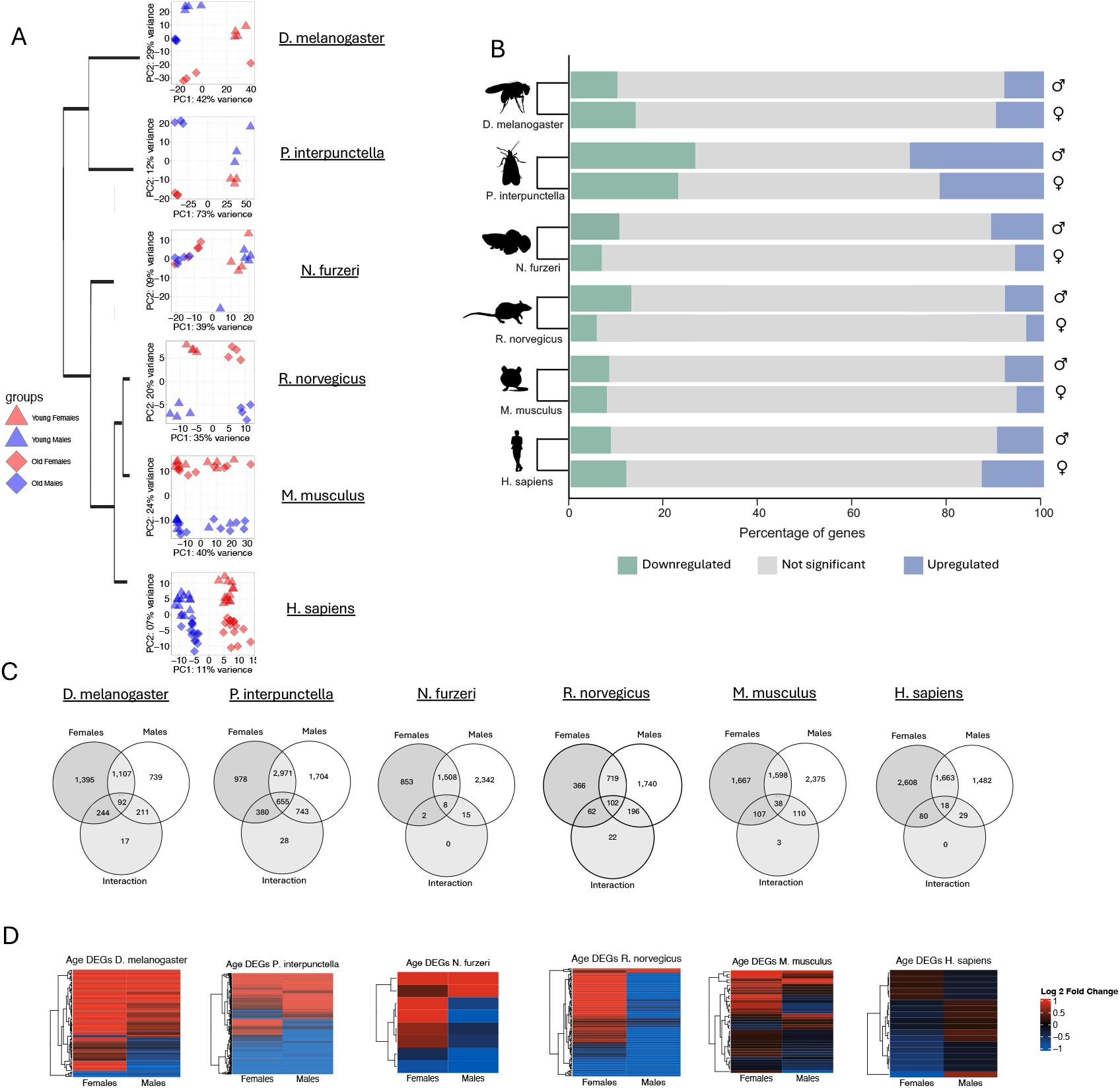
Principal Component Analysis (PCA) and differentially express gene (DEG) breakdown for each species. A) PCA plots for each species in the same order as the phylogenetic tree from figure 1A. Points within PCA plots are colored by sex (female = red, male = blue), with shapes denoting age (young = triangle, old = diamond). B) Percentage of DEGs in age for each species and sex. Bars are colored according to significance and regulation direction (downregulated = green, not significant = grey, upregulated = blue). C) Venn diagrams of all DEGs in age for each species. Numbers reflect all DEGs which areunique to that comparison. D) Heatmap of the regulation DEGs that were shared between females, males, and interaction (the center most number in the Venn diagrams). Colors correspond to the log2 fold change of each gene in age with red > 0 and blue < 0. **Alt text.** Figures highlighting con-specific RNA-seq results.

### Age-Associated Transcriptional Change Between the Sexes Varied Among Species

To examine how aging alters sex-specific transcription within each species, we identified age-associated differentially expressed genes (agDEGs) using three contrasts: female old vs. young, male old vs. young, and the age-by-sex interaction. Across species, females exhibited anywhere from 1,249-4,984 agDEGs, while males exhibited anywhere from 2,149-6,073 agDEGs (**Table 1; Supplemental File 1**), indicating substantial species variation, including sample size, power, and variance that affect the magnitude of age-associated transcriptional changes (**Supplemental Fig. 1**). Additionally, the direction of transcriptional change varied between sexes, within species (**Fig. 2B**, **Table 1**). In females, *D. melanogaster, N. furzeri,* and *M. musculus* showed a significant bias (Chi-square test, *p <* 0.05) toward upregulated agDEGs, whereas *R. norvegicus* and *H. sapiens* exhibited a predominance of downregulated agDEGs (Chi-square test, *p <* 0.05), and *P. interpunctella* showed no significant directional bias. In males, *D. melanogaster* was the only species with a significant (Chi-square test, *p <* 0.05) enrichment of upregulated agDEGs, while *P. interpunctella, N. furzeri, R. norvegicus*, and *H. sapiens* showed a higher proportion of downregulated agDEGs (Chi-square test, *p <* 0.05), and *M. musculus* showed no significant bias. These patterns highlight substantial diversity in both the magnitude and direction of transcriptional aging responses across evolutionary lineages.

**Table 1.** Number of differentially expressed genes for each species based on each contrast.

| Species | contrast | upregulated | downregulated | Total DEGs |
| --- | --- | --- | --- | --- |
| *D. melanogaster* | Male Old - Male Young | 1,146 | 1,003 | 2,149 |
|  | Female Old - Female Young | 1,639 | 1,199 | 2,838 |
|  | Age x Sex Interaction | 117 | 447 | 564 |
| *P. interpunctella* | Male Old - Male Young | 2,951 | 3,122 | 6,073 |
|  | Female Old - Female Young | 2,526 | 2,458 | 4,984 |
|  | Age x Sex Interaction | 850 | 956 | 1,806 |
| *N. furzeri* | Male Old - Male Young | 1,862 | 2,011 | 3,873 |
|  | Female Old - Female Young | 1,246 | 1,125 | 2,371 |
|  | Age x Sex Interaction | 10 | 15 | 25 |
| *R. norvegicus* | Male Old - Male Young | 1,056 | 1,701 | 2,757 |
|  | Female Old - Female Young | 585 | 664 | 1,249 |
|  | Age x Sex Interaction | 57 | 325 | 382 |
| *M. musculus* | Male Old - Male Young | 2,085 | 2,036 | 4,121 |
|  | Female Old - Female Young | 1,954 | 1,456 | 3,410 |
|  | Age x Sex Interaction | 98 | 160 | 258 |
| *H. sapiens* | Male Old - Male Young | 1,502 | 1,690 | 3,192 |
|  | Female Old - Female Young | 2,077 | 2,292 | 4,369 |
|  | Age x Sex Interaction | 89 | 38 | 127 |

To assess sex differences in aging, we quantified the proportion of agDEGs that were female-specific, male-specific, or shared between sexes within each species (**Fig. 2C**; **Supplemental File 2**). Interestingly, across species, 12-44% of agDEGs were female-specific, while 19-54% were male-specific across species. The proportion of common agDEGs ranged from 25-48%, with *P. interpunctella* exhibited the greatest overlap while *R. norvegicus* showed the least overlap (**Fig. 2C**; **Supplemental File 2**). Together, these results demonstrate that muscle aging processes are associated with widespread transcriptional changes that are highly variable across species and sex, in both magnitude and regulatory direction.

To further investigate sex differences in aging, it is important to identify the genes that show age-associated transcriptional trajectories that differ between the sexes. Thus, we evaluated DEGs identified from age-by-sex interaction effects (absDEGs) and compared them with agDEGs identified within each sex (**Fig. 2C, Supplemental File 3**). Interestingly, absDEGs that overlapped with shared agDEGs often exhibited differences in the magnitude of expression change between sexes within a given species. In some cases, these genes showed opposing regulatory trajectories, with age-associated upregulation in one sex and downregulation in the other (**Fig. 2D**). For example, in *H. sapiens*, *TPRG1*, with its antisense transcript *TPRG1-AS1* are implicated in cardiovascular aging (Ren et al 2022), exhibited age-related upregulation in males but downregulation in females (**Fig. 2D)**, suggesting a potential link to the higher incidence of cardiovascular disease observed in males (Freedman et al 2026). These patterns indicate that sex differences in aging are not necessarily driven by entirely distinct gene sets, but rather by quantitative and directional differences in specific gene regulation.

Together, these findings demonstrate that aging induces widespread yet highly variable transcriptional changes across species and sexes. While some species exhibit strong concordance between males and females, others display pronounced sex-biased aging signatures, reinforcing the importance of incorporating both sexes in comparative analyses of aging.

### The Regulation of Canonical Human Aging Genes Differed Across Species

To gain functional insight into agDEGs and absDEGs, we classified all genes into orthogroups, which are sets of genes descended from a single ancestral gene that may include both orthologs and paralogs across species (Emms and Kelly 2019). Among six species, 95.2% of all annotated genes were assigned successfully to 23,024 orthogroups containing one or more species using OrthoFinder2 (**Supplemental File 4**), with only 8% of these orthogroups being classified as species-specific, and the remaining 4.8% of annotated genes being unassigned. This result indicates that the vast majority of genes belong to conserved gene families shared across at least two species. Overall, 5,343 orthogroups contained representatives from all six species, reflecting substantial evolutionary conservation. However, only 449 orthogroups were strict single-copy (1:1) orthologs (**Supplemental File 5**), highlighting the limited prevalence of highly conserved, non-duplicated gene families across these phylogenetically distant taxa. Together, these orthogroup assignments provided the comparative framework for downstream functional and comparative analyses of aging-associated genes.

To assess whether well-characterized human aging genes exhibit shared transcriptional responses across species, we compared 307 aging-associated genes from the GenAge database (de Magalhães et al 2024) with the set of 1:1 orthologs identified in our dataset. Only eight genes overlapped across all six species: *CDK7, CISD2, ERCC2, GCLM, MLH1, PARP1, STUB1,* and *XPA* (**Supplemental Fig. 2**). Among these, *CISD2* exhibited one of the most consistent signatures. *CISD2* localizes to the endoplasmic reticulum and mitochondrial membranes and plays a role in cytosolic Ca²⁺ homeostasis (Shen et al 2021). We observed significant age-related downregulation of *CISD2* in *M. musculus, H. sapiens,* and *P. interpunctella*, suggesting a shared decline in *CISD2* expression during aging across both vertebrates and insects. This finding is consistent with prior studies linking reduced *CISD2* expression to mitochondrial dysfunction and accelerated tissue aging (Chen et al 2009).

The remaining species exhibited a mixture of expression patterns among these eight shared human aging genes, reflecting lineage-specific divergence in canonical aging regulators. Although these genes are well-established in human aging biology, their transcriptional trajectories vary substantially across species and between sexes, highlighting both shared elements and species-specific adaptations. Moreover, only four of the eight genes were identified as downregulated agDEGs in human skeletal muscle, which may be due to tissue specific expression patterns as many genes cataloged in the human aging database were derived from brain or other non-muscle tissues. This highlights the importance of tissue context when interpreting shared aging signatures.

Furthermore, the relatively small number of shared 1:1 orthologs limited the number of previously defined aging genes that could be compared directly across all species. This limitation highlights the need for more inclusive comparative frameworks, such as incorporating paralogs within broader orthogroups, to better capture the full spectrum of shared and divergent aging mechanisms across species. Together, these findings emphasize the importance of comparative analyses for interpreting the evolutionarily shared aging pathways and caution against assuming that transcriptional aging signatures identified in a single species, a single sex, or a single tissue type readily generalize across taxa and biological contexts.

### Evolutionarily Shared Aging-Associated Orthogroups were Identified Across All Species

To compare patterns of orthogroup-level aging signatures across species, sexes, and lineages, we generated an UpSet plot (**Fig. 3A; Supplemental File 6**), which highlighted key intersection sets of orthogroups containing agDEGs. Notably, three orthogroups contained at least one agDEG from both sexes from all six species, representing the most shared transcriptional aging signatures identified in this study (**Fig. 3A**). At broader phylogenetic scales, the amount of shared orthogroups increased in closely related species. For instance, there was a greater amount of shared orthogroups (N= 27) when the rodents were grouped together (*R. norvegicus* and *M. musculus*) then there were when all the mammals were grouped together (N= 6, Rodents and *H. sapiens*). This indicates a negative relationship between evolutionary distance and conservation of transcriptional aging signatures.

**Figure 3.**
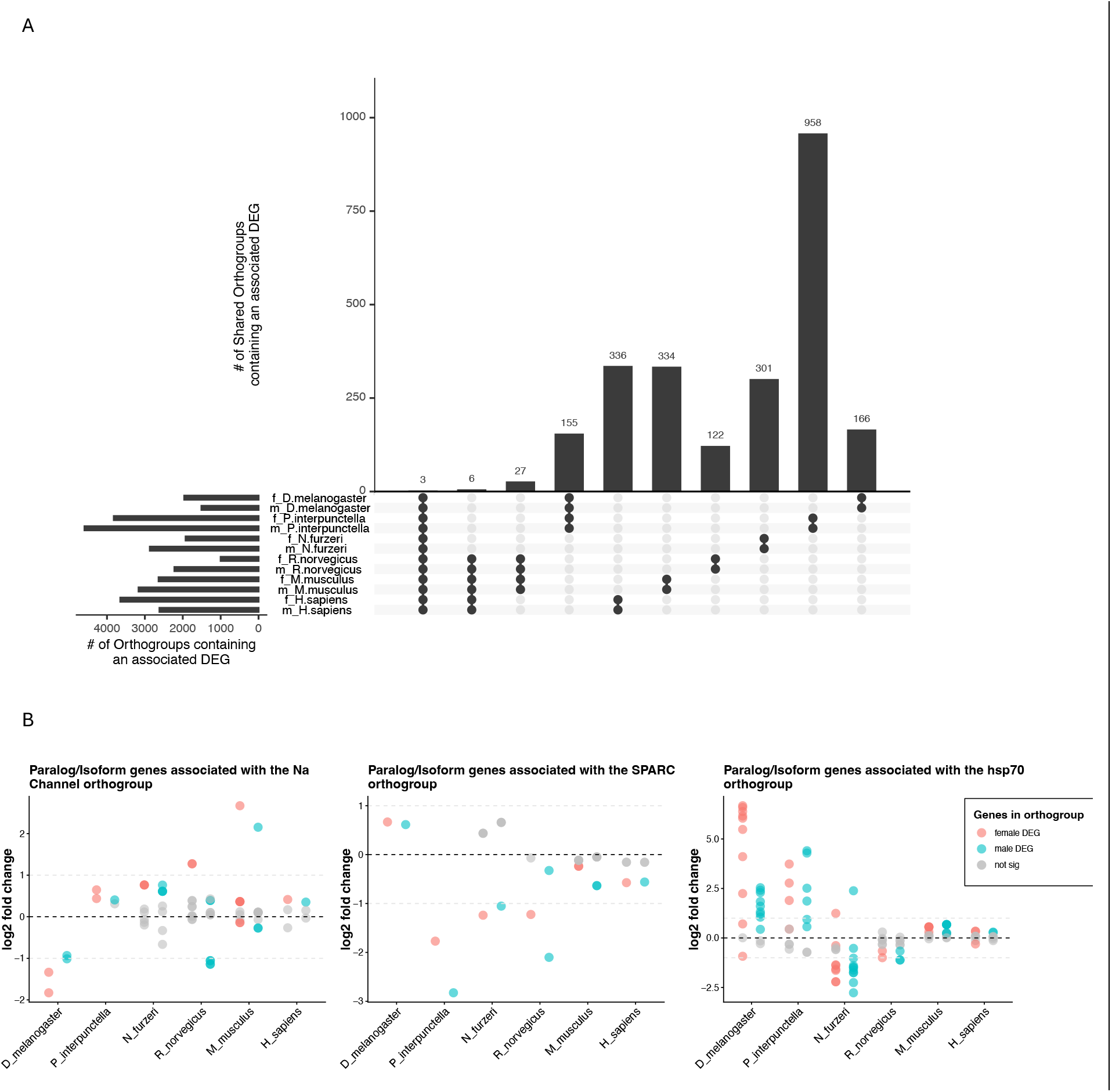
A) An upset plot of the full dataset highlighting the three shared orthogroups. Upset plot of both sexes and all species illustrating the amount of orthogroups common or unique to different combinations of samples. The first bar is the three shared orthogroups for which both sexes of each species contributed an agDEG. The next bar is mammal specific, followed by rodent specific, insect specific, and then the six species specifics. B) Shared orthogroups. These graphs depict the log fold change in age of all genes within the orthogroup from both sexes and all six species. Colors represent sex and significance with red = female differentially expressed genes (DEGs), blue = male DEGs, grey = nonsignificant genes. **Alt text.** Figures showing amounts of orthogroups selected species and sexes contributed to and plots of the three shared orthogroups.

The three orthogroups with agDEGs in males and females of all six species (**Fig. 3B**) included genes that encode Na⁺ channel proteins, heat shock protein 70 (HSP70), and secreted protein acidic and cysteine rich (SPARC). Across species and sexes, these orthogroups showed significant age-associated transcriptional changes. The SPARC orthogroup was predominantly downregulated with age in all species except *D. melanogaster*, whereas the other two shared orthogroups exhibited greater variability across species. We further examined age-associated expression patterns within the HSP70 orthogroup (**Fig. 3B**), a key regulator of proteostasis and a well-established hallmark of aging (López-Otín et al 2023). Across the phylogeny, both up- and downregulation were observed, and the dominant direction and magnitude of change varied among species and sexes. Insects (*D. melanogaster* and *P. interpunctella*) showed the strongest age-associated upregulation, particularly in female *D. melanogaster,* whereas *M. musculus* and *H. sapiens* exhibited more moderate expression increases. In contrast, *N. furzeri* and *R. norvegicus* displayed a higher proportion of downregulated *HSP70* agDEGs, with greater magnitude observed in *N. furzeri* (**Fig. 3B; Supplemental File 7**). Together, these results indicate that HSP70 expression change is a shared feature of aging, even though the direction, magnitude, and gene-specific contributions vary substantially across species and sexes.

### Lineage-Specific Expression Shifts in HSP70 Reveal Divergent Evolution of Aging Regulation

To investigate the evolutionary dynamics of gene expression within the HSP70 orthogroup, we applied an OU model along the phylogeny of orthologous HSP70 genes across the six species. This framework identifies branches where observed expression levels deviate significantly from evolutionary expectations (Rohlfs et al 2013), here defined as expression shifts, representing lineage-specific changes in gene regulation within homologous gene families. Such shifts provide insight into how genes diverge in regulation over evolutionary time, including how the expression of paralogous genes diverge.

Multiple expression shifts with distinct directions and magnitudes were detected across the HSP70 orthogroup (**Fig. 4A**). The most pronounced shift was associated with speciation between the *D. melanogaster* HSP70 and *P. interpunctella* HSP70 orthogroups. In *P. interpunctella,* a gene duplication related expression shift was observed among LOC*128682120, LOC128680252* and *LOC128670490*, with LOC*128682120* exhibiting low expression and a lack of aging-related upregulation in females. Within *D. melanogaster,* HSP70 paralogous genes (*HSP68, HSP70BA, HSP70BB, HSP70BC, HSP70BBB, HSP70AA,* and *HSP70AB*) were strongly differentially expressed with age in both sexes, with most exhibiting significant age-by-sex interaction effects and pronounced age-associated upregulation in females (**Fig. 4B**). An exception was *D. melanogaster* paralog *HSP70BA*, which lacked this age-by-sex interaction but instead showed a distinct expression shift toward overall expression reduction relative to other clade members (**Fig. 4A-B**), suggesting that this overall low expression may underlie the loss of sex-age interaction.

**Figure 4.**
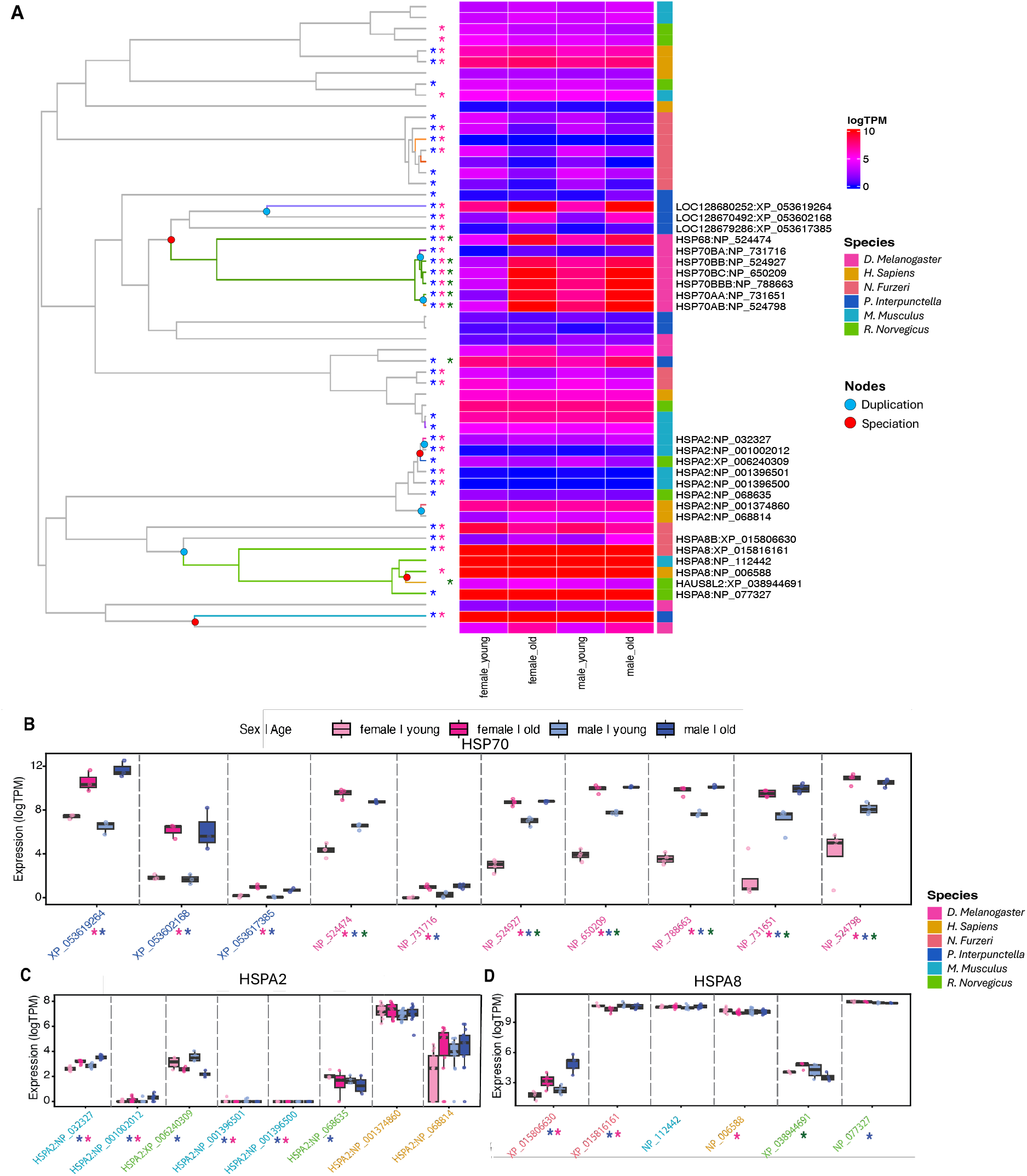
Application of the Ornstein–Uhlenbeck (OU) model to the HSP70 orthogroup. A) Expression shifts were detected on specific branches of the gene tree within this orthogroup and are highlighted in different colors, representing the direction and magnitude of lineage-specific expression shifts. The heatmap displays log-transformed TPM (logTPM) values across four traits: female young, female old, male young, and male old. The color bar on the right indicates the species origin of each gene. Asterisks denote differentially expressed genes (DEGs): blue indicates male-specific DEGs, pink indicates female-specific DEGs, and green indicates genes showing significant age-by-sex interaction effects. Gene symbols concatenated with protein IDs are shown. B–D) Boxplots showing expression levels of selected genes from the HSP70 family: HSP70 genes B), HSPA8 genes C), and HSPA2 genes D) across the four groups (female young, female old, male young, and male old). Colors of the protein IDs indicate the species of origin. Asterisks denote the same DEG categories as described in panel A). **Alt text.** Figures demonstrating expression shifts detected using an OU model.

A second major expression shift associated with gene duplication events was identified within the vertebrate HSPA8 clade, including *N. furzeri* (*HSPA8* and *HSPA8B*), *M. musculus*, *H. sapiens*, and *R. norvegicus* (*HSPA8* and *HAUS8L2*) (**Fig. 4A**). In *N. furzeri*, the paralog *HSPA8B* exhibited significantly lower overall expression relative to *HSPA8*. Similarly, *HAUS8L2*, a paralog of *HSPA8* in *R. norvegicus*, showed a lineage-specific shift toward reduced expression compared to *HSPA8* orthologs in rats and other vertebrates (**Fig. 4A and 4C**). Interestingly, unlike the age-associated downregulation observed for *HSPA8*, both *N. furzeri HSPA8B* and *R. norvegicus HAUS8L2* displayed an inverted sex-biased aging pattern, with age-associated upregulation in females but downregulation in males, a pattern not observed among other vertebrate *HSPA8* orthologs (**Fig. 4C**).

Additional lineage-specific expression shifts associated with both speciation and gene duplication were identified within the HSPA2 orthogroup among mammals, including *M. musculus* (NP_032327 vs. NP_001002012), *R. norvegicus* (XP_006240309 vs. NP_068635), and *H. sapiens* (NP_001374860 vs. NP_068814) (**Fig. 4A**, **4D**). Overall, the human *HSPA2* lineage exhibited a speciation-associated shift toward elevated baseline expression relative to rodents. Interestingly, *HSPA2* was classified as an agDEG in both rodent species but not in humans (**Fig. 4D**), suggesting divergence in the magnitude of age-associated regulation across species. Within each species, paralog pairs also displayed distinct expression shifts, indicating regulatory divergence following duplication events (**Fig. 4D**). Across mammals, *M. musculus* and *H. sapiens HSPA2* orthologs showed age-associated upregulation in both sexes, whereas *R. norvegicus* exhibited age-associated downregulation, further supporting species-specific regulation within this conserved HSP70 chaperone family.

Together, these results demonstrate that while HSP70-family genes are implicated broadly in aging, the direction, and magnitude of expression shifts associated with speciation and/or duplication vary across evolutionary lineages. These findings highlight both the shared involvement of proteostasis pathways in aging and the lineage-specific regulatory divergence that shapes how these pathways are modulated across species.

### Sex-Biased Orthogroups Reveal Divergent Muscle Aging Pathways

To identify sex-dependent patterns of aging at the orthogroup level, we analyzed agDEG contributions separately for males and females across species. We defined sex-biased orthogroups as those containing at least one agDEG from all species of one sex, but not consistently represented across all species of the opposite sex. This definition captures gene families exhibiting shared sex-biased transcriptional responses to aging across taxa. Using these criteria, we identified four female-biased and 18 male-biased age-associated orthogroups (**Fig. 5A-B; Supplemental Fig. 3-4; Supplemental File 8**). Similar to the globally shared aging signatures, the degree of orthogroup conservation decreased with increasing evolutionary distance, indicating that sex-biased aging programs become progressively more lineage-specific across phylogeny.

**Figure 5.**
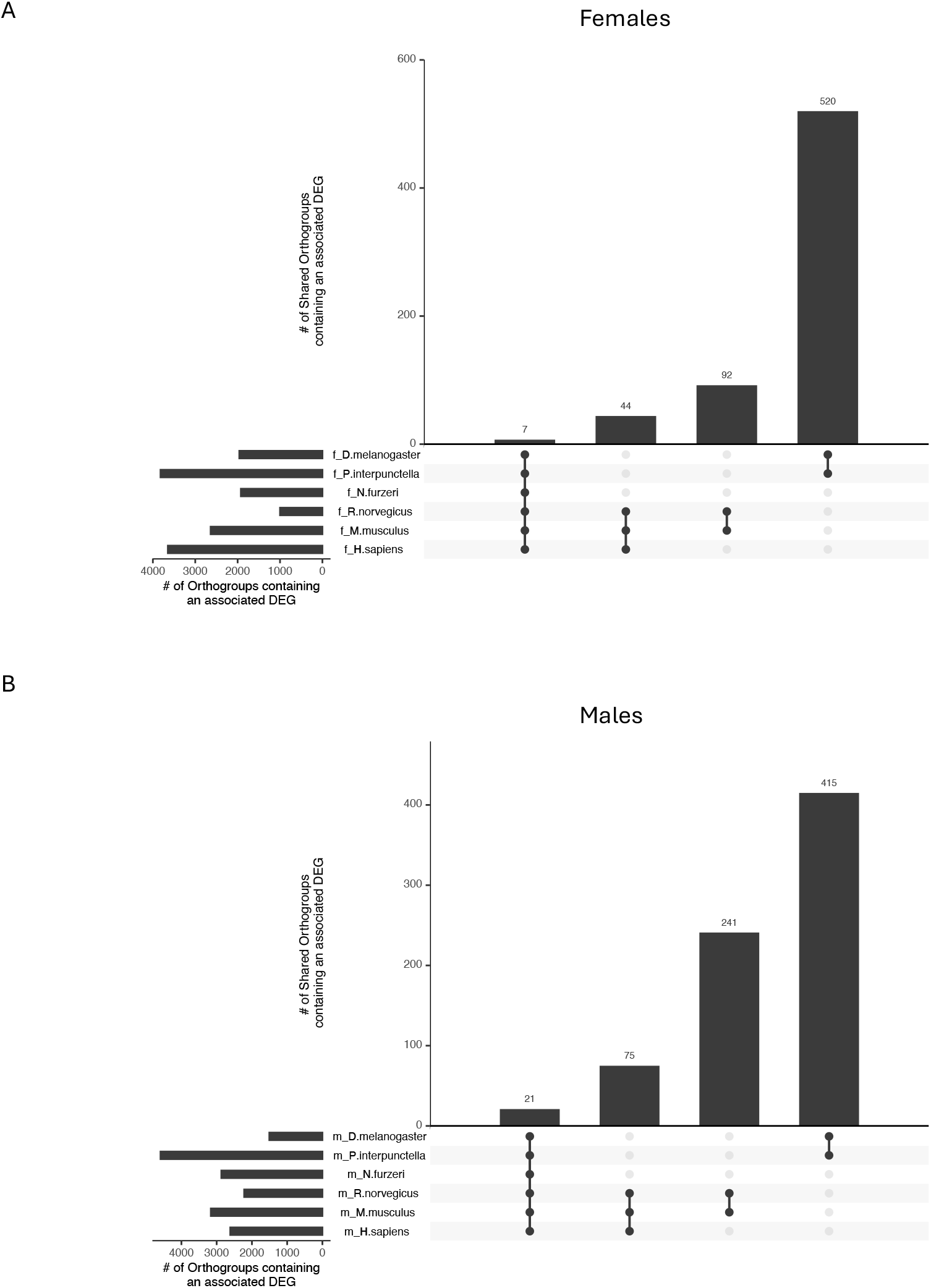
Sex-biased aging markers found in the transcription data. Upset plots indicating the amount of orthogroups (number on top of bars) that each species contributed to with an agDEG. A) are females only and B) are males only. The first bars of each upsetplot are orthogroups for which all species contributed to. The next bars are mammals only, followed by insects only, and rodents only. **Alt text.** Figures highlighting the amount of sex-biased orthogroups that select species contributed to.

Sex-biased orthogroups reflected unique aging patterns between the sexes. For example, the four female-biased orthogroups were dual-specificity tyrosine-regulated kinase (Dyrk), phosphodiesterase, kinase heavy chain, and collagens, all of which exhibited diverse female-biased age-associated expression changes (**Fig. 6A and Supplemental Fig. 3**). These genes are involved in muscle differentiation and health, autophagy, as well as muscle contraction and muscle stiffness. In contrast, male-biased orthogroups were enriched for genes associated with mitochondrial function and cellular metabolism (**Fig. 6B and Supplemental Fig. 4**), including voltage-dependent anion channels *(VDAC1–3)*, acetyl-CoA metabolism *(ACSS)*, and aminolevulinate synthase *(ALAS1)*. Genes involved in ATP production and muscle energetics included V-type proton ATPase components, as well as Na⁺/K⁺-ATPase subunits *(ATP1B1)*.

**Figure 6.**
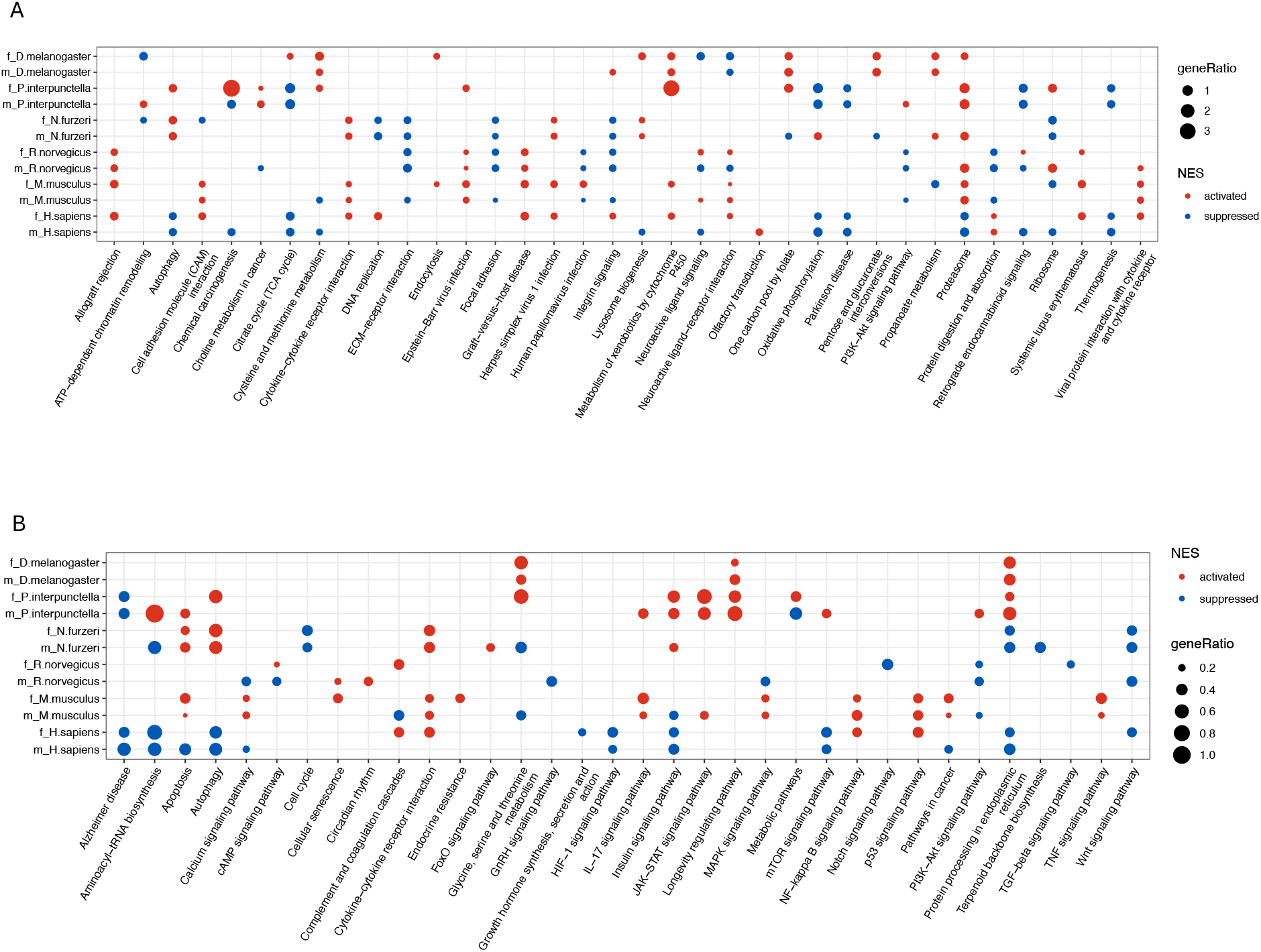
KEGG pathway comparison between species. A) are the two most significant activated and the two most significant suppressed KEGG pathways in age from both sexes of all species. B) KEGG pathways that were cross-referenced and subset to include only those pathways that were also listed as common aging pathways in the Aging Atlas online database. For both plots, color of dots indicate whether the pathway is activated (red) or suppressed (blue) in age based on its normalized enrichment score (NES) from the gene set enrichment analysis (GSEA). **Alt text.** Figures comparing the activation/suppression/no-change of KEGG pathways in age across all species and sexes.

Although the direction and magnitude of age-associated regulation varied among species, these orthogroups were represented consistently across males but showed less consistent involvement in females. These patterns suggest that age-associated differences in mitochondrial and metabolism pathways are conserved more strongly in males, whereas females show stronger conservation of pathways related to functional muscle decline. Together, these results demonstrate that sex-biased aging signatures are not driven by isolated genes, but rather by coordinated, functionally coherent orthogroups, highlighting distinct evolutionary trajectories in how males and females regulate key physiological systems during aging.

### Pathways of Cellular Stress and Metabolism Decline During Aging

To identify the biological processes most affected by aging across species and sexes, Kyoto Encyclopedia of Genes and Genomes (KEGG) pathway enrichment analyses were carried out for each species and sex. Despite the substantial phylogenetic diversity of the dataset, several core aging-associated pathways emerged consistently as the most significantly enriched across taxa (**Fig. 6A**). For example, pathways involved in proteostasis, including proteasome, protein digestion and absorption, and autophagy, all were enriched significantly in multiple species. These pathways mediate cellular responses to age-associated stress, such as the accumulation of misfolded proteins, oxidative damage, and impaired protein turnover (Kourtis and Tavernarakis 2011). As central components of protein quality control, these systems typically are challenged during aging, reflecting an increased cellular stress burden (Higuchi-Sanabria et al 2018).

Additionally, when KEGG pathways were cross-referenced to known aging pathways from the Aging Atlas online database (Aging Atlas Consortium 2020), additional pathways relating to proteostasis were identified such as protein processing in endoplasmic reticulum (**Fig. 6B**), further reflecting the increase of cellular damage in age within the majority of the species studied. A full list of all enriched KEGG pathways for both sexes in each species can be found in **Supplemental File 10**.

Interestingly, the direction of age-associated expression changes differed across species. In *H. sapiens*, proteostasis-related pathways, including autophagy, proteasome, and protein processing in endoplasmic reticulum were expressed lower in muscles with age (**Fig. 6A-B)**, suggesting a reduced capacity for cellular proteostasis proteasome activities (Saez and Vilchez 2014, Njomen and Tepe 2019) and consistent with increased proteolysis in human muscle aging (Ubaida-Mohien et al 2019). In contrast, many non-human species, such as *N. furzeri, M. musculus,* and *R. norvegicus,* exhibited increased expression in some of these pathways, consistent with a compensatory mechanism to to increased proteotoxic stress (Morimoto and Cuervo 2014, Hamazaki and Murata 2024). These contrasting patterns highlight both shared involvement and divergent regulation of cellular stress response mechanisms during aging.

In addition to proteostasis pathways, several metabolic and structural processes were enriched broadly with age across species (**Fig. 6**). These included thermogenesis and pathways associated with muscle fiber organization and cytoskeletal integrity. The decreased expression in pathways of oxidative phosphorylation, PI3K-Akt signaling, calcium signaling, and muscle contraction pathways is consistent with well-established declines in skeletal muscle function, metabolic flexibility, and mitochondrial efficiency during aging (Chen et al 2021). This pattern was not witnessed in all species, or in both sexes within species, demonstrating sex-specific and evolutionary divergent patterns of enrichment.

Comparative analysis across taxa revealed that the most consistently enriched pathways involve proteostasis, reinforcing its central role as a core feature in aging biology. At the same time, the presence of sex-biased and lineage-specific enrichment patterns highlights evolutionary divergence in how organisms respond to cellular stress and allocate resources during aging.

Together, these findings indicate that aging is characterized by a shared increase in cellular stress, while the transcriptional strategies used to manage this process diverges across species and between sexes.

## Discussion

This study provides a cross-species analysis of sex-biased transcriptional aging across diverse animal lineages. We identified three aging markers shared by all species and both sexes, including proteostasis-related genes such as *HSP70*, highlighting the evolutionary importance of this core aging process. Aging signatures were predominantly sex-biased, defined as genes or pathways that were significantly altered with age in the same sex of all species but not consistently represented in the other sex across all species This is in contrast to sex-specific signatures, which would entail being represented in only one sex across all species and never in the other sex. These signatures relayed involvement of structural and functional properties in female-biased agDEGs and mitochondrial and metabolic properties in male-biased agDEGs. In parallel, we observed substantial lineage-specific divergence, with decreasing overlap of aging-associated orthogroups across evolutionary distances and lineage-dependent expression shifts in key gene families. Functionally, we identified shared proteostasis disruption and male-biased regulation in mitochondrial functions, demonstrating that aging reflects both shared molecular mechanisms and divergent regulatory strategies across species and sexes.

Our study advances the field of aging biology by moving beyond single-species analyses to a comparative, cross-species framework that explicitly incorporates sex as a central biological variable. While previous transcriptomic studies have largely focused on a limited set of model organisms in mice (Jin et al 2025, Bilgic et al 2025), flies (Lu et al 2023, Sang et al 2025), and worms (Wang et al 2022, Ham et al 2022), our approach integrates both model and non-model species and includes both sexes, enabling a systematic evaluation of shared and evolutionary divergent sex-biased aging-associated transcriptional programs.

Importantly, this study demonstrates that the biology of aging cannot be described fully from single-species or sex-biased perspectives. Instead, it emerges as a comprehensive process shaped by shared cellular stress responses and modulated by lineage specification and sex-specific physiology. By integrating comparative transcriptomics within an evolutionary framework, our work provides a scalable approach for identifying robust, cross-species aging biomarkers, and offers new insights into how fundamental aging processes are both shared and diversified across the animal kingdom.

### Shared Molecular Hallmarks and Divergent Regulatory Strategies of Muscle Aging

We were able to identify three shared orthogroups (Na⁺ channel proteins, HSP70, and SPARC) demonstrating that similar core transcriptional features of aging are present across deeply divergent species and both sexes. Importantly, these orthogroups map to fundamental muscle biological function, including cellular structural integrity (SPARC), muscle membrane excitability (Na⁺ channels), and proteostasis (HSP70), suggesting that aging consistently impacts these core cellular functions in muscle. While the manner in which these orthogroups are altered with age differs between species, and sometimes between the sexes, their identification provides an exciting opportunity to explore these nuances in future aging studies.

One can begin to parse out some of these nuances in orthogroup regulation between species by investigating the fold change of agDEGs within these shared orthogroups. For example, *SPARC* exhibited consistent age-associated downregulation across species, with the exception of *D. melanogaster*. SPARC is a matricellular glycoprotein strongly associated with muscle fibrosis, degeneration, and sarcopenia (Ghanemi et al 2022a), and its decline during aging is consistent with previous observations in mammalian muscle aging (Gruber et al 2005, Ghanemi et al 2022b). In contrast, *D. melanogaster* showed increased *SPARC* expression, potentially associated with the tissue-specific extracellular matrix remodeling and fibrosis-like accumulation during aging (Vaughan et al 2018). Similarly, Na⁺ channel genes (*SCN*) were predominantly upregulated across species, again with the exception of *D. melanogaster*. This pattern may represent a compensatory response to maintain cellular excitability in the context of declining Na⁺/K⁺-ATPase (NKA) activity during aging (Wyckelsma and McKenna 2016).

The HSP70 orthogroup is a particularly strong candidate for a shared aging mechanism because HSP70 proteins are central to maintaining proteostasis. The change in HSP70 expression with age across all species supports the idea that proteostasis efficiency is declining, or at the very least being altered with age, consistent with previous findings (López-Otín et al 2023). The observation of both upregulation and downregulation of agDEGs within the HSP70 orthogroup within a given species likely reflects distinct strategies of proteostatic responses. Increased expression of HSP70 genes may represent a compensatory response to accumulating cellular stress, as HSP70 proteins function as molecular chaperones that protect against elevated oxidative stress and chronic inflammation, as observed in aged *D. melanogaster* (Tower 2010) and during *H. sapiens* muscle aging (Cumming et al 2021). On the other hand, decreased *HSP70* expression may indicate a progressive decline in proteostasis capacity, resulting in reduced ability to manage misfolded proteins and increased susceptibility to cellular damage, including muscle atrophy and impaired regeneration, as reported in skeletal muscle of aged *R. norvegicus* (Vasilaki et al 2002) and elderly *H. sapiens* (Senf 2013). These variations suggest that different species may employ distinct transcriptional mechanisms to fine-tune proteostasis during aging.

Together, these findings highlight that aging involves shared disruption of core biological systems, while the regulatory responses to these disruptions diverge across species and sexes, reflecting evolutionary differences in how organisms cope with cellular stress and functional decline.

### Sex-Biased Muscle Aging Reflects Divergent Regulation of Collagens and Mitochondrial Metabolism

The identification of sex-biased orthogroups across species establishes that aging is not a uniform process, but is driven instead by sex-dependent regulatory programs. These molecular signatures arise from coordinated, pathway-level changes, revealing fundamentally distinct strategies by which females and males undergo aging. Our comparative framework further demonstrates that these sex-biased patterns are shared across evolutionary lineages, rather than being species-specific phenomena. For instance, the distinct separation of transcriptomic profiles by sex in five of the six species highlights that sex can be a dominant driver of aging-associated transcriptional variation. While some aging-related pathways, such as those involved in proteostasis (HSP70), are shared across sexes, others exhibit clear sex-bias, for example, the consistent involvement of muscle degeneration pathways in females and mitochondrial and metabolic pathways in males.

Consistent with this divergence, aging in females was characterized by enrichment of orthogroups associated with sarcopenia and structural components such as collagens, suggesting the degeneration of normal muscle functioning in age. The downregulation of agDEGs of *DYRK* in females of all species, except *P. interpunctella*, could be an indicator of muscle degeneration as some paralogs of *DYRK* have been linked to sarcopenia (Jiao et al 2025). Interestingly enough, the consistent downregulation of the agDEGs phosphodiesterase (*PDE4* specifically) in females of all species, except *N. furzeri*, is suggestive of a compensatory mechanism to combat muscle degeneration, as inhibition of *PDE4* has been identified as a therapeutic target in sarcopenia drug treatments (Balasubramaniam et al 2018, Angelino et al 2025). Finally, the variable expression patterns between species of agDEGs of kinase heavy chain and collagen in all females relays species specific alterations in muscle function and tissue stability with age respectively (Holwerda et al 2025). Together, these findings support a model in which aging in females involves a multi-level perturbation of muscle homeostasis, ultimately impacting muscle function.

In contrast, male-biased agDEG orthogroups were enriched for genes involved in mitochondrial function and cellular metabolism, including voltage-dependent anion channels (*VDAC*) and enzymes involved in acetyl-CoA metabolism and redox regulation, indicating age-associated mitochondrial functional decline in aged males (Viña et al 2003, Camus et al 2012, Demarest and McCarthy 2014). Additional male agDEGs associated with ATP production and ion homeostasis, such as creatine kinases, V-type proton ATPase components, and Na⁺/K⁺-ATPase subunits, further indicate alterations in energy production and cellular energetics during aging (Wyckelsma and McKenna 2016). These changes directly correlate with diminished ATP replenishment, reduced metabolic rate, and impaired ion homeostasis (Alves et al 2023). These patterns suggest that aging in males is characterized by a more pronounced disruption of mitochondrial function and metabolic homeostasis, which are essential for sustaining muscle performance.

Despite these distinct molecular trajectories, both sexes converge on a similar physiological outcome, age-associated decline in muscle function (Trombetti et al 2015). However, the underlying mechanisms differ: females exhibit more alterations in muscle structure, whereas males show greater alterations in mitochondrial and metabolic pathways. This divergence highlights that similar aging phenotypes can arise from different molecular regulatory mechanisms, shaped by sex-specific physiology.

Together, this work extends our current understanding of aging by the systematic analysis of transcriptomes from both young and old individuals of both sexes from six very diverse species. Until recently, aging studies have been male-biased, which reflected a long-standing bias in biomedical research (Beery and Zucker 2011). The presence of distinct, shared molecular pathways underlying aging in each sex suggests that interventions targeting age-related decline may need to be sex-specific, as the efficacy of lifespan extending interventions often differs between males and females (Hajdarovic et al 2026). More broadly, this work highlights the importance of incorporating both sexes in comparative and translational studies to more accurately capture the complexity of aging biology.

### Orthogroup-Based Comparative Framework Captures Evolutionary Complexity of Muscle Aging

A key methodological strength of this study is the use of orthogroup-based analysis, rather than relying solely on 1:1 ortholog matching across taxa. Although 1:1 orthologs are used commonly to infer direct evolutionary relationships and functional equivalence (Altenhoff et al 2012), this approach excludes duplicated or lineage-expanded genes, which often play critical roles in adaptation and species-specific biology (Koonin 2005). Moreover, the assumption of strict functional equivalence among 1:1 orthologs increasingly has been questioned (Studer and Robinson-Rechavi 2009, Stamboulian et al 2020). Furthermore, while the number of 1:1 orthologs is relatively high in closely related species, the number decreases substantially with greater evolutionary divergence, limiting their application in broad comparative analyses such as ours.

By contrast, orthogroups, including gene families descended from a common ancestral gene, enable a more comprehensive representation of evolutionary relationships. Using this framework, we analyzed 23,024 orthogroups across six species, compared to only 449 1:1 orthologs, substantially expanding the scope of cross-species comparison. This approach allowed us to identify shared functional modules while accommodating lineage-specific gene expansions and losses (Gabaldón and Koonin 2018), which are particularly relevant in the context of aging. Although orthogroup-based analyses may introduce some ambiguity in precise functional equivalence, they provide a more integrative and evolutionarily informative view of transcriptomic divergence (Sheng et al 2022). Future studies combining this approach with functional validation, such as protein modeling or experimental validation, will be important to resolve gene-specific roles within complex orthogroups.

### Conclusions and Future Directions

This study reveals that muscle aging is governed by a combination of shared and sex-biased molecular programs that operate across vast evolutionary distances. The consistent age-associated expression change of proteostasis-associated HSP70 orthogroups across taxa provides compelling evidence that cellular stress response and proteostasis remodeling represent universal features of aging (Haigis and Yankner 2010). At the same time, our analyses highlight sex divergence in aging, with the consistent involvement of mitochondrial and metabolic pathways in males, demonstrating that aging trajectories can diverge systematically between sexes even within shared biological systems. Our integrative framework, combining comparative transcriptomics, orthogroup-based evolutionary mapping, and OU modeling, establishes a scalable and evolutionarily informed strategy for dissecting the molecular architecture of aging beyond traditional model organisms. By moving beyond single-gene comparisons to pathway- and orthogroup-level analyses, this approach captures both shared functional modules and lineage-specific regulatory divergence, providing a more comprehensive view of aging biology. As transcriptomic resources expand, non-model species will enable increasingly powerful comparative analyses, allowing researchers to disentangle how sex, ecology, and life history strategies shape the evolution of aging. Ultimately, these insights will be critical for advancing a more generalizable and mechanistically grounded understanding of aging, with direct implications for the development of sex-informed and evolutionarily robust therapeutic anti-aging strategies.

## Material/Methods

### Animal Sample Collection

*D. melanogaster* (DGRP 304 strain, Bloomington Drosophila Stock Center, RRID: BDSC_25177) were reared at the University of Alabama at Birmingham in incubators with 12 hours light/12 hours dark conditions at 25 °C in 65% humidity and fed Jazz-Mix Drosophila Food (Fisher), supplemented with active dry yeast. Virgin flies from both sexes were collected over the course of 48 hours and housed in single sex groups of 20 in separate vials, totaling about 15 vials per sex, used for aging. For the young group, at 8 days post eclosion, flies were flash frozen in liquid nitrogen, and thoraxes were dissected on a dry ice-cooled surface. Five to ten thoraxes were pooled for each sample (n = 4) for both sexes to be used in RNA extraction. The same procedure was repeated at the old time point (54 days post eclosion). Total RNA was extracted using the Monarch Total RNA miniprep kit (New England Biolabs) following the manufacturer’s protocol.

*P. interpunctella* were raised from pools of larvae housed in 32 oz. plastic cups, held at 25 °C in 14 hours light/10 hours dark conditions, and fed a custom mix of cracked wheat, wheat bran, wheat germ, yeast, honey, glycerin, and water (Silhacek and Murphy 2008). When larvae reached the fifth larval instar, they were transferred to individual 1 oz. deli cups for the remainder of their lifespan. Young samples were collected on the day they eclosed, while old samples were collected at 13 days post eclosion. The cup housing the animal to be collected was placed on ice for 5-15 minutes until the animal stopped moving. The body was then segmented into head, thorax, and abdomen. Individual thoraxes were placed into 1.5 mL microcentrifuge tubes containing 300 μL TRIzol (Invitrogen, ThermoFisher, US) and stored at -80 °C. Total RNA was extracted using TRIzol following the manufacturer’s protocol.

### RNA Sequencing

For both insect samples described above, 400ng of total RNA was submitted to Novogene (Sacramento, CA, USA) for library preparation and sequencing. Briefly, RNA concentration and integrity were assessed using Qbit and Bioanalyzer. High-quality RNA was poly-A selected and reverse transcribed using random hexamer primers. cDNA libraries were prepared by Novogene and were sequenced on an Illumina NovaSeq platform, generating on average 40 million paired-end 150 base-pair reads per sample.

### Public Dataset Curation

RNA-seq datasets of four species were obtained from NCBI public datasets (**Table 2**). Raw reads were retrieved from the Sequence Read Archive (SRA) database. Details of each dataset, including replicate numbers (**Fig. 1B**), accession numbers, and experimental metadata, are provided in **Table 2**.

**Table 2.**
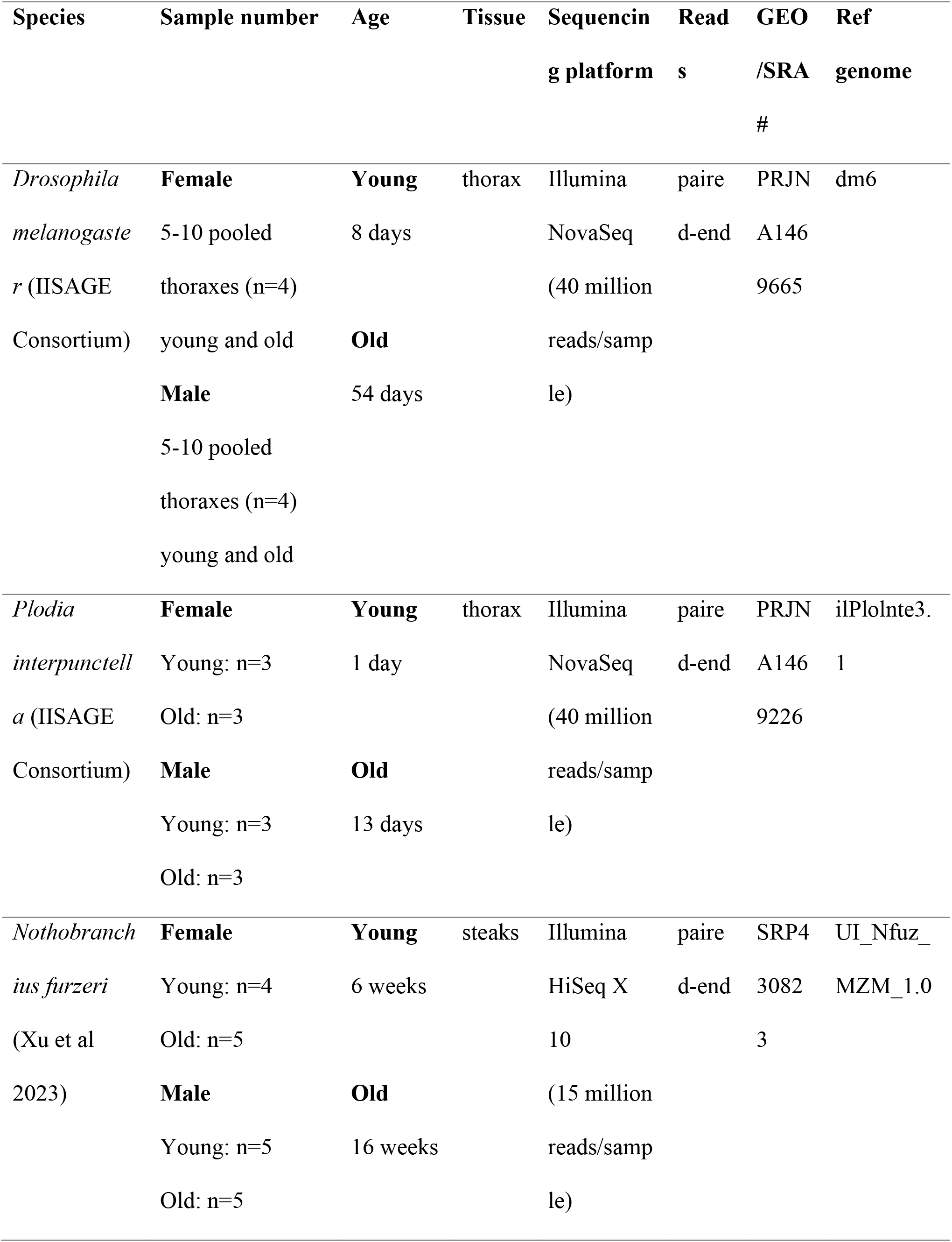
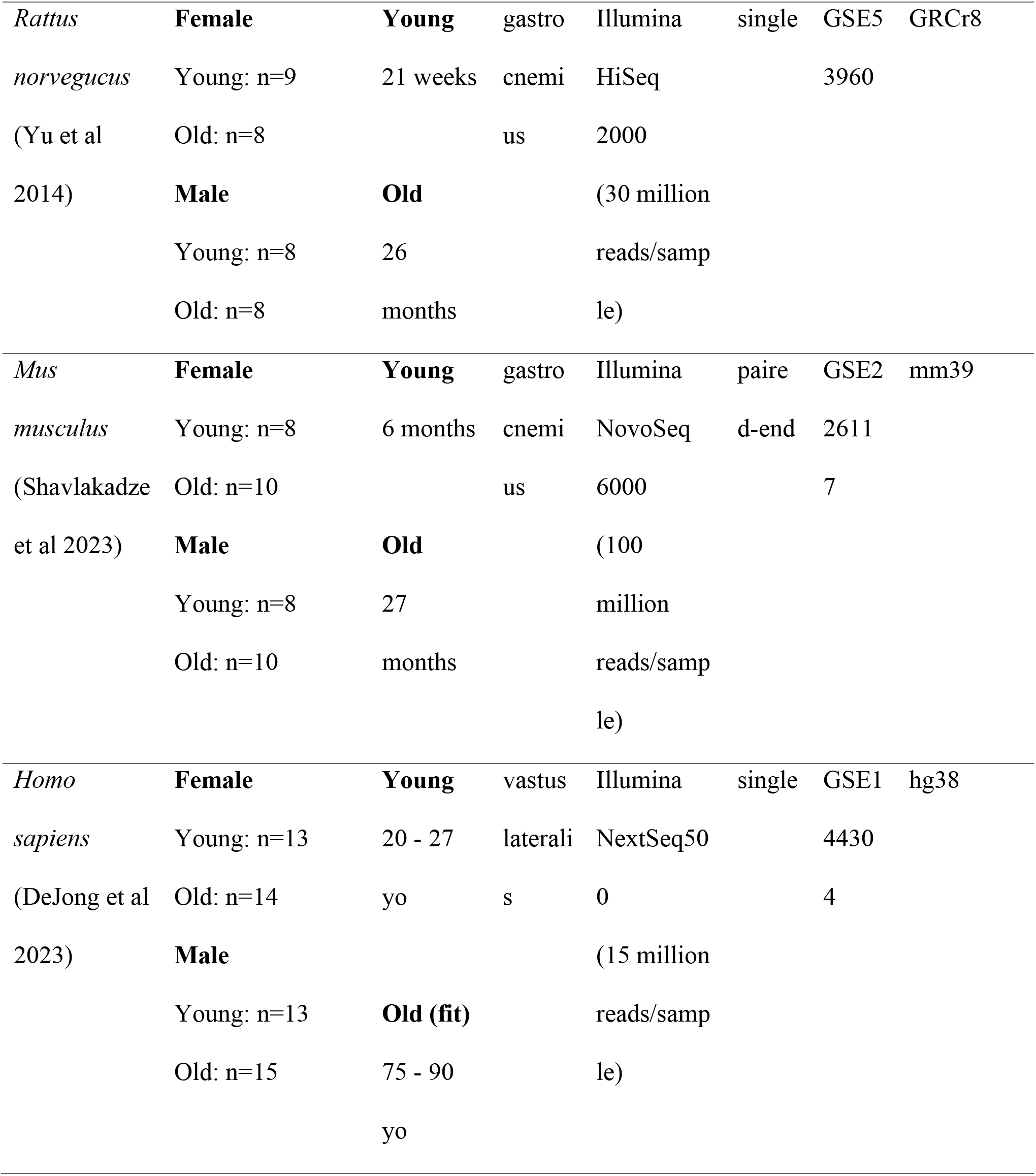
Sample specifics used in the workflow.

All samples were from bulk RNA-seq studies that included both sexes and at least two age groups representing young and old timepoints. For species with multiple timepoints, such as *M. musculus* (Shavlakaez et al 2023) and *R. norvegicus* (Yu et al 2014), only the youngest sexually mature and the oldest available age group were included to represent young and old. In the *R. norvegicus* dataset containing technical duplicates (Yu et al 2014), no significant differences were observed between these duplicates. Therefore, the corresponding raw reads were concatenated before processing. We note that sequencing strategies varied across studies. For example, *R. norvegicus* (Yu at al 2014) and *H. sapiens* (DeJong et al 2023) studies used single-read sequencing, while all the other studies used paired-end sequencing. Library preparation methods also differed: ribo-depleted libraries were used for *R. norvegicus* (Yu et al 2014), *M. musculus* (Shavlakadze et al 2023), and *N. furzeri* (Xu et al 2023), while poly-A selected libraries were used for the remaining species. All studies included had RNA-seq data derived from skeletal muscle tissue, with sampling sites as follows: vastus lateralis (*H. sapiens*), gastrocnemius (*R. norvegicus* and *M. musculus*), and steak (*N. furzeri*).

### Differential Gene Expression

Raw reads were processed using kallisto (Bary et al 2016) for species-specific pseudoalignment and transcript quantification. Reference genomes obtained from NCBI (**Table 2**) included *D. melanogaster* (Release 6 plus ISO1 MT, dm6), *P interpunctella* (ilPlolnte3.1 alternate haplotype), *N. furzeri* (UI_Nfuz_MZM_1.0), *R. norvegicus* (GRCr8), *M. musculus* (CRCm39, mm39), and *H. sapiens* (GRCh38.p14, hg38). Transcript quantifications for all species were imported into the statistical environment R (R Core Team 2023) using the tximport package (Soneson et al 2015). Transcript abundances were summarized to the gene level and converted to counts using the tximport option “tx2gene” and a generated TxDB transcript to gene object generated from the reference Gene Transfer Format files (gtf) from each species using the R package GenomicFeatures (Lawrence et al 2013). Differentially expressed genes in age (agDEGs) were identified for each species using DESeq2 (Love et al 2014). These agDEGs were identified within each sex, and additional contrasts were performed to detect age-by-sex interaction effects. Genes were considered differentially expressed based on an adjusted p-value (p.adjust) < 0.05. Differences in the proportion of up-versus downregulated agDEGs for both sexes of each species were then investigated using a Chi-Squared Goodness of Fit Test.

### Cross-species Orthogroup Determination

Orthogroups were identified using the online tool OrthoFinder2 (Emms and Kelly 2019), with the complete proteome of each species obtained from Ensembl (release version 110; Dyer et al 2025) as input. To account for isoforms, or multiple transcripts from the same gene, only the longest (primary) transcript variant from each gene was used for this analysis. OrthoFinder2 was run with its default settings, which performed all-verse-all BLAST searches and used sequence similarity scores to construct gene trees and identify orthogroups.

OrthoFinder2 outputs a tab-separated values file (**Supplemental File 10**) and arbitrarily names each orthogroup. Then, the names of the orthogroups were changed to reflect the proteins that make up the orthogroup. The protein identifier names for each species within an orthogroup were searched in NCBI, and the consensus protein name was used to name the orthogroup.

Next, species-specific agDEGs were mapped to their encoded proteins and assigned to corresponding orthogroups. Orthogroups containing agDEGs from two or more species were classified as common aging markers. Species-specific agDEG contributions to shared orthogroups were visualized using the UpSet R package (Gehlenborg 2019). For universal and sex-biased common orthogroups, the median log2 fold-change (old:young) across species and sexes was plotted using ggplot2 (Wickham 2016). An illustration of the complete bioinformatic workflow used in this manuscript can be found in **Fig. 1C**.

### Functional Enrichment Analysis

Species-specific gene functional annotation databases were constructed for *N. furzeri* and *P. interpunctella* using AnnotationForge (Carlson and Pagès 2023). Gene Set Enrichment Analysis (GSEA) was performed using clusterProfiler (Wu et al 2021, Yu et al 2012) on the full set of results from each contrast for each species/sex combination to identify enriched KEGG pathways. As *P. interpunctella* was not listed in the KEGG database, their KEGG pathways were obtained using GhostKOALA (Kanehisa et al 2016) and incorporated into the GSEA analysis in clusterProfiler. Enriched KEGG pathways were then compared across all species and sexes to identify shared and sex-biased functional patterns of aging.

### Detection of Expression Shifts Using The Ornstein-Uhlenbeck Model

To detect evolutionary shifts in agDEG expression, OU models were applied to orthogroup gene trees generated by OrthoFinder. For each orthogroup, mean log-TPM values were calculated for each age-sex group combination (young-female; young-male; old-female; old-male), which were treated as multivariate traits. Expression evolution was modeled as a multivariate model using the R package l1ou (v1.43), which identifies branch-specific shifts in expression optima (θ) along a phylogeny. Conceptually, this approach compares evolutionary models with and without allowing changes in optimal expression levels on specific branches and determines whether introducing such shifts provides a better explanation of the observed expression patterns across species.

Model comparison was performed using the Akaike Information Criterion corrected for small sample size (AICc), a widely used metric that balances model fit and model complexity. A branch was inferred to have an expression shift when allowing a change in expression optimum on that branch resulted in a better-supported evolutionary model than one assuming a single expression optimum across the tree.

We applied the l1ou framework following its standard assumptions (Khabbazian et al 2016), which uses the Ornstein-Uhlenbeck process along a phylogenetic tree to provide functions to study trait evolution from comparative transcriptomics and detect past changes in the expected mean gene expression values. The strength of stabilizing selection (α) and the variance of the stochastic component (σ²) were assumed to be constant across branches, yielding a single global stationary variance (γ). Expression levels across four sex-age groups were modeled as correlated multivariate traits, with α and σ² estimated jointly. Shifts in expression optima were assumed to occur along the same branches across traits.

## Data availability

All scripts for this project can be found at this link:

https://github.com/ecr77/Defining_Ubiquitous_and_Sex-

**Biased_Aging_Biomarkers_Using_Transcriptional_Data_across_Evolutionar…**

## Acknowledgements -Funding

This work was funded by the National Science Foundation Grant 2213824 to the IISAGE symposium. We would also like to express our appreciation to Heidi Johnson, MS, for collecting the *D. melanogaster* samples.

## Author contributions

E.C.R. curated publicly available RNA-seq datasets for four of the species, and performed the cross-species transcriptomic analyses to identify agDEGs. Y.F. contributed to RNA-seq dataset collection, data analysis discussions, bioinformatics pipeline construction and conducted the OU model analyses and result interpretation. C.K. collected *P. interpunctella* samples for RNA sequencing. R.P.M., J.W., T.G., G.W., E.L., A.W., P.R.B., N.C.R., and J.E.D. contributed to result interpretation and discussion. E.C.R. and Y.F. drafted the original manuscript and designed the figures under the supervision of J.E.D., P.R.B., and N.C.R. All authors involved reviewed, revised, and approved the final manuscript.

## Conflict of Interest

The authors declare that they have no competing interests.

## Notes

### Competing Interest Statement

The authors have declared no competing interest.

## References

Aging Atlas Consortium. 2020. Aging Atlas: a multi-omics database for aging biology. Nucleic Acids Res. gkaa894 doi:10.1093/nar/gkaa894

Altenhoff AM, Studer RA, Robinson-Rechavi M, Dessimoz C. 2012. Resolving the ortholog conjecture: Orthologs tend to be weakly, but significantly, more similar in function than paralogs. PLoS Comput. Biol. 8(5):e1002514 doi:10.1371/journal.pcbi.1002514

Alves FM, Ayton S, Bush AI, Lynch GS, Koopman R. 2023. Age-related changes in skeletal muscle iron homeostasis. J. Gerontol. A. Biol. Sci. Med. Sci. 78(1):16–24 doi:10.1093/gerona/glac139

Angelino E, Bodo L, Sartori R, Malacarne V, D’Anna B, Formaggio N, Barua S, Raiteri T, Lauria A, Reano S, Murabito A, Nicolau M, Ferrero F, Pezzini C, Rossino G, Favero F, Valmasoni M, Filigheddu N, Menga A, Corà D, Hirsch E, Olivero S, Sartorelli V, Proserpio V, Ghigo A, Sandri M, Porporato PE, Talarico D, Caretti G, Graziani A. 2025. Nat. Metab. 7:2548–2570 doi:10.1038/s42255-025-01397-5

Augustin H, Partridge L. 2009. Invertebrate models of age-related muscle degeneration. Biochim. Biophys. Acta – Gen. Subj. 1790(10):1084–1094 doi:10.1016/j.bbagen.2009.06.011

Balasubramaniam A, Sheriff S, Friend A, James JH. 2018. Phosphodiesterase 4B knockout prevents skeletal muscle atrophy in rats with brain injury. Am. J. Physiol. Regul. Integr. Comp. Physiol. 315(2):R417–R421 doi:10.1152/ajpregu.00042.2018

Ballantyne AJ, Rogers WA. 2011. Sex bias in studies selected for clinical guidelines. J. Women’s Health 20(9):1297–1306 doi:10.1089/jwh.2010.2604

Barr MM, García LR, Portman DS. 2018. Sexual dimorphism and sex differences in *Caenorhabditis elegans* neuronal development and behavior. Genetics 208(3):909–935 doi: 10.1534/genetics.117.300294

Bary NL, Pimentel H, Melsted P, Pachter L. 2016. Near-optimal probabilistic RNA-seq quantification. Nat. Biotechnol. 34:525–527 dio:10.1038/nbt.3519

Beery AK, Zucker I. 2011. Sex bias in neuroscience and biomedical research. Neurosci. Biobehav. Rev. 35(3):565–572 doi:10.1016/j.neubiorev.2010.07.002

Biga PR, Duan JE, Young TE, Marks JR, Bronikowski A, Decena LP, Randolph EC, Pavuluri AG, Li G, Fang Y, Wilkinson GS, Singh G, Nigrin NT, Larschan EN, Lonski AJ, Riddle NC, IISAGE Consortium. 2025. Hallmarks of aging: a user’s guide for comparative biologists. Aging Res. Rev. 104:102616 doi:10.1016/j.arr.2024.102616

Bilgic M, Obata R, Panfil VI, Zhu Z, Saeki M, Gotoh Y, Kishi Y. 2025. Age-associated transcriptomic and epigenetic alterations in mouse hippocampus. Aging Cell 24(11):e70233 doi:10.1111/acel.70233

Bronikowski AM, Meisel RP, Biga PR, Walters JR, Mank JE, Larschan E, Wilkinson GS, Valenzuela N, Conard AM, de Magalhães JP, Duan JE, Elias AE, Gamble T, Graze RM, Gribble KE, Kreiling JA, Riddle NC. 2022. Sex-specific aging in animals: Perspective and future directions. Aging Cell 21:e13542 doi: 10.1111/acel.13542

Camus MF, Clancy DJ, Dowling DK. 2012. Mitochondria, maternal inheritance, and male aging. Curr. Biol. 22(18):P1717–1721 doi:10.1016/j.cub.2012.07.018

Carlson M, Pagès H. 2023. AnnotationForge: Tools for building SQLite-based annotation data packages. R package version 1.40.2 <https://nioconductor.org/packages/AnnotationForge>

Carmody C, Duesing GG, Kane AE, Mitchell SJ. 2022. Is sex as a biological variable still being ignored in preclinical aging research? J. Gerontol. A Biol. Sci. Med. Sci. 77(11):2177–2180 doi:10.1093/gerona/glac042

Chen X, Li M, Chen B, Wang W, Zhang L, Ji Y, Chen Z, Ni X, Shen Y, Sun H. 2021. Transcriptome sequencing and analysis reveals the molecular mechanism of skeletal muscle atrophy induced by denervation. Ann. Transl. Med. 9(8):697 doi:10.21037/atm-21-1230

Chen YF, Kao CH, Chen TY, Wang CH, Wu CY, Tsai CY, Liu FC, Yang CW, Wei YH, Hsu MT, Tsai SF, Tsai TF. 2009. Cisd2 deficiency drives premature aging and causes mitochondria-mediated defects in mice. Genes & Dev. 23:1183–1194 doi:10.1101/gad.1779509

Critchlow AJ, Alexander SE, Hiam DS, Ferrucci L, Scott D, Lamon S. 2025. Associations between female sex hormones and skeletal muscle aging: The Baltimore longitudinal study of aging. J. Cachexia Sarcopenia Muscle 16:e13786 doi:10.1002/jcsm.13786

Cumming KT, Kyamme NH, Schaad L, Ugelstad I, Raastad T. 2021. Muscular HSP70 content is higher in elderly compared to young, but is normalized after 12 weeks of strength training. Eur. J. Appl. Physiol. 121(6):1689–1699 doi:10.1007/s00421-021-04633-4

[dataset]* DeJong JCBC, Attema BJ, van der Hoek MD, Verschuren L, Caspers MPM, Kleemann R, van der Leij FR, van der Hoek AM, Nieuwenhuizen AG, Keijer J. 2023. Sex differences in skeletal muscle-aging trajectory: same processes, but with different ranking. GeroScience 45:2367–2386 dio: 10.1007/s11357-023-00750-4

de Magalhaes JP, Abidi Z, Dos Santos GA, Avelar RA, Barardo D, Chatsirisupachai K, Clark P, De-Souza EA, Johnson EJ, Lopes I, Novoa G, Senez L, Talay A, Thornton D, To PKP. 2024. Human aging genomic resources: updates on key databases in aging research. Nucleic Acids Res. 52(D1):D900–D908 doi10.1093/nar/gkad927

Demarest TG, McCarthy MM. 2014. Sex differences in mitochondrial (dys)function: Implications for neuroprotection. J. Bioenerg. Biomembr. 47(1-2):173–188 doi:10.1007/s10863-014-9583-7

Dyer S, Austine-Orimoloye O, Azov AG, Barba M, Barnes I, Barrera-Enriquez VP, Becker A, Bennett R, Beracochea M, Berry A, Bhai J, Bhurji SK, Boddu S, Lins PRB, Brooks L, Ramaraju SB, Campbell LI, Martinez MC, Charkhchi M, Cortes LA, Davidson C, Denni S, Dodiya K, Donaldson S, Houdaigui BE, Naboulsi TE, Falola O, Fatima R, Genez T, Martinez JG, Gurbich T, Hardy M, Hollis Z, Hunt T, Kay M, Kaykala V, Lemos D, Lodha D, Mathlouthi N, Merino GA, Merritt R, Mirabueno LP, Mushtaq A, Hossain SN, Pérez-Silva JG, Perry M, Piližota I, Poppleton D, Prosovetskaia I, Raj S, Salam AIA, Saraf S, Saraiva-Agostinho N, Sinha S, Sipos B, Sitnik V, Steed E, Suner MM, Surapaneni L, Sutinen K, Tricomi FF, Tsang I, Urbina-Gómez D, Veidenberg A, Walsh TA, Willhoft NL, Allen J, Alvarez-Jarreta J, Chakiachvili M, Cheema J,da Rocha JB, De Silva NH, Giorgetti S, Haggerty L, Ilsley GR, Keatley J, Loveland JE, Moore B, Mudge JM, Naamati G, Tate J, Trevanion SJ, Winterbottom A, Flint B, Frankish A, Hunt SE, Finn RD, Freeberg MA, Harrison PW, Martin FJ, Yates AD. 2025. Ensembl 2025. Nucleic Acids Res. 53(D1):D948–D957 doi:10.1093/nar/gkae1071

Emms DM, Kelly S. 2019. OrthoFinder: phylogenetic orthology inference for comparative genomics. Genome Biol. 20:238 dio:10.1186/s13059-019-1832-y

Freedman AA, Colangelo LA, Ning H, Borrowman JD, Lewis CE, Schreiner PJ, Khan SS, Lloyd-Jones DM. 2026. Sex differences in age of onset of premature cardiovascular disease and subtypes: the coronary artery risk development in young adults study. J. Am. Heart Assoc. 15(3):e044922 doi:10.1161/JAHA.125.044922

Gabaldón T, Koonin EV. 2013. Functional and evolutionary implications of gene orthology. Nat. Rev. Genet. 14(5):360–366 doi:10.1038/nrg3456

Gehlenborg N. 2019. UpSetR: A more scalable alternative to Venn and Euler Diagrams for visualizing intersecting sets. R package version 1.4.0 <https://CRAN.R-project.org/package=UpSetR>

Ghanemi A, Yoshioka M, St-Amand J. 2022a. Genetic expression between aging and exercise: secreted protein acidic and rich in cysteine as a potential “exercise substitute” antiageing therapy. Genes (Basel) 13(6):950 doi:10.3390/genes13060950

Ghanemi A, Yoshioka M, St-Amand J. 2022b. Secreted protein acidic and rich in cysteine as an exercise-induced gene: towards novel molecular therapies for immobilization-related muscle atrophy in elderly patients. Genes (Basel) 13(6):1014 doi:10.3390/genes13061014

Gruber HE, Sage EH, Norton HJ, Funk S, Ingram J, Hanley Jr EN. 2005. Targeting deletion of the SPARC gene accelerates disc degeneration in the aging mouse. J. Histochem. Cytochem. 53(9):1131–1138 doi:10.1369/jhc.5A6687.2005

Haigis MC, Yankner BA. 2010. The aging stress response. Mol. Cell 40:333–344 doi:10.1016/j.molcel.2010.10.002

Hajdarovic KH, Riddle NC, Webb AE. 2026. Sex differences in response to longevity interventions. Aging Res. Rev. 118:103123 doi:10.1016/j.arr.2026.103123

Ham S, Kim SS, Park S, Kim EJE, Kwon S, Park HEH, Jung Y, Lee SJV. 2022. Systematic transcriptome analysis associated with physiological and chronological aging in *Caenorhabditis elegans*. Genome Res. 32(11-12):2003–2014 doi:10.1101/gr.276515.121

Hamazaki J, Murata S. 2024. Relationship between protein degradation, cellular senescence, and organismal aging. J. Biochem. 175(5):473–480 doi:10.1093/jb/mvae016

Higuchi-Sanabria R, Frankino PA, Paul III JW, Tronnes SU, Dillin A. 2018. A futile battle? Protein quality control and the stress of aging. Dev. Cell 44(2):139–163

Holdcroft A. 2007. Gender bias in research: now does it affect evidence based medicine? J R Soc Med. 100(1):2–3 doi:10.1177/014107680710000102

Holwerda AM, Aussieker T, Senden J, Bouwman FG, van Kranenburg J, Overman A, Snijders T, Verdijk LB, Holm L, van Loon LJC. 2025. Assessing the composition of myofibrillar and muscle connective protein fractions within skeletal muscle tissue. Crit. Rev. Food Sci. Nutr. 66(1):248–258 doi:10.1080/10408398.2025.2522363

Human aging genomic resources: updates on key databases in aging research. Nucleic Acids Res. 52(D1):D900–D908

Iliadi KG, Iliadi NN, Boulianne GL. 2009. Regulation of *Drosophila* life-span: effect of genetic background, sex, mating and social status. Exp. Gerontol. 44(8):546–553 doi: 10.1016/j.exger.2009.05.008

Jiao XF, Mu GJ, Zhoa WY, Ni R, Zhao C, Lu X, Wu JQ, Gao W, Luo L. 2025. Dyrk1b as a potential biomarker for sarcopenia in older adults. BMC Geriatrics 25:278 doi:10.1186/s12877-025-05942-5

Jin K, van Velthoven CTJ, Kaplan ES, Glattfelder K, Barlow ST, Boyer G, Carey D, Casper T, Chakka AB, Chakrabarty R, Clark M, Departee M, Desierto M, Gary A, Gloe J, Goldy J, Guilford N, Guzman J, Hirschstein D, Lee C, Liang E, Pham T, Reding M, Ronellenfitch K, Ruiz A, Sevigny J, Shapovalova N, Shulga L, Sulc J, Torkelson A, Tung H, Levi B, Sunkin SM, Dee N, Esposito L, Smith KA, Tasic B, Zeng H. 2025. Brain-wide cell-type-specific transcriptomic signatures of healthy aging in mice. Nature 638:182–196 doi:10.1038/s41586-024-08350-8

Kanehisa M, Sato Y, Morishima K. 2016. BlastKOALA and GhostKOALA: KEGG tools for functional characterization of genome and metagenome sequences. J. Mol. Biol. 428(4):726–731 dio:10.1016/j.jmb.2015.11.006

Khabbazian M, Kriebel R, Rohe K, Ane C. 2016. Fast and accurate detection of evolutionary shifts in Ornstein-Uhlenbeck models. Methods Ecol. Evol. 7(7):811–824 doi:10.1111/2041-210X.12534

Koonin EV. 2005. Orthologs, paralogs, and evolutionary genomics. Annu. Rev. Genet. 39:309–338 doi:10.1146/annurev.genet.39.073003.114725

Kourtis N, Tavernarakis N. 2011. Cellular stress response pathways and ageing: intricate molecular relationships. EMBO J. 30(13):2520–2531 doi:10.1038/emboj.2011.162

Lemaître JF, Ronget V, Tidiėre M, Allainė D, Berger V, Cohas A, Colchero F, Conde DA, Garratt M, Liker A, Marais GAB, Scheuerlein A, Szėkely T, Gaillard JM. 2020. Sex differences in adult lifespan and aging rates of mortality across wild mammals. PNAS 117(15):8546–8553 doi:10.1073/pnas.1911999117

López-Otín C, Blasco MA, Partridge L, Serrano M, Kroemer G. 2023. Hallmarks of aging: An expanded universe. Cell 186(2):P243–278 doi:10.1016/j.cell.2022.11.001

Love MI, Huber W, Anders S. 2014. Moderated estimation of fold change and dispersion for RNA-seq data with DESeq2. Genome Biol. 15(12):550 dio:10.1186/s13059-014-0550-8

Lu TC, Brebić M, Park YJ, Jackson T, Chen J, Kolluru SS, Qi Y, Katheder NS, Cai XT, Lee S, Chen YC, Auld N, Liang CY, Ding SH, Welsch D, D’Souza S, Pisco AO, Jones RC, Leskovec J, Lai EC, Bellen HJ, Luo L, Jasper H, Quake SR, Li H. 2023. Aging Fly Cell Atlas identifies exhaustive aging features at cellular resolution. Sci. 380(6650):eadg0934 doi:10.1126/science.adg0934

Marais GAB, Gaillard JM, Vieira C, Plotton I, Sanlaville D, Gueyffier F, Lemaitre JF. 2018. Sex gap in aging and longevity: can sex chromosomes play a role? Biol. Sex Differ. 9:33 doi:10.1186/s13293-018-0181-y

McCulloch D, Gems D. 2003. Evolution of male longevity bias in nematodes. Aging Cell 2:165–173 dio: 10.1046/j.1474-9728.2003.00047.x

Mendoza E, Moen DS, Holt NC. 2023. The importance of comparative physiology: mechanisms, diversity and adaptation in skeletal muscle physiology and mechanics. J. Exp. Biol. 226:jeb245158 doi:10.1242/jeb.245158

Millington JW, Riedout EJ. 2018. Sex differences in *Drosophila* development and physiology. Curr. Opin. Physiol. 6:46–56 doi: 10.1016/j.cophys.2018.04.002

Morimoto RI, Cuervo AM. 2014. Proteostasis and the aging proteome in health and disease. J. Gerontol. A Biol. Sci. Med. Sci. 69(Suppl 1):S33–S38 doi:10.1093/gerona/glu049

Mukund K, Subramaniam S. 2019. Skeletal muscle: A review of molecular structure and function, in health and disease. WIREs Syst. Biol. Med. 12:e1462 doi:10.1002/wsbm.1462

Njomen E, Tepe JJ. 2019. Proteasome activation as a new therapeutic approach to target proteotoxic disorders. J. Med. Chem. 62(14):6469–6481 doi:10.1021/acs.jmedchem.9b00101

R Core Team. 2023. _R: a language and environment for statistical computing_. R Foundation for Statistical Computing, Vienna Austria. <https://www.R-project.org/>

Reichard M, Blažek R, Žåk J, Cellerino A, Polačik M. 2012. The sources of sex differences in aging in annual fish. J. Anim. Ecol. 91:540–550 dio:10.1111/1365-2656.13656

Ren X, Zhu H, Deng K, Ning X, Li L, Liu D, Yang B, Shen C, Wang X, Wu N, Chen S, Gu D, Wang L. 2022. Long noncoding RNA TPRG1-AS1 suppresses migration of vascular smooth muscle cells and attenuates atherogenesis via interacting with MYH9 protein. Arterioscler. Thromb. Vasc. Biol. 42(11):1378–1397 doi:10.1161/ATVBAHA.122.318158

Rohlfs RV, Harrigan P, Nielsen R. 2013. Modeling gene expression evolution with an extended Ornstein-Uhlenbeck process accounting for within-species variation. Mol. Biol. Evol. 31(1):201–211 doi:10.1093/molbev/mst190

Saez I, Vilchez D. 2014. The mechanistic links between proteasome activity, aging and age-related diseases. Curr. Genomics 15(1):38–51 doi:10.2174/138920291501140306113344

Sang Y, Ning X, Xu Q, Wang L, Yan Y, Zhang L, Bi X. 2025. Characterization of transcriptomics during aging and genes required for lifespan in *Drosophila* intestine. Sci. Rep. 15:14695 doi:10.1038/s41598-025-98888-y

Senf SM. 2013. Skeletal muscle heat shock protein 70: diverse functions and therapeutic potential for wasting disorders. Front. Physiol. 4:330 doi:10.3389/fphys.2013.00330

[dataset]* Shavlakadze T, Xiong K, Mishra S, McEwen C, Gadi A, Wakai M, Salmon H, Stec MJ, Negron N, Ni M, Wei Y, Atwal GS, Bai Y, Glass DJ. 2023. Age-related gene expression signatures from limb skeletal muscles and the diaphragm in mice and rats reveal common and species-specific changes. Skelet. Muscle 13:11 dio:10.1186/s13395-023-00321-3

Shen ZQ, Huang YL, Teng YC, Wang TW, Kao CH, Yeh CH, Tsai TF. 2021. CISD2 maintains cellular homeostasis. BBA – Mol. Cell Res. 1868(4):118954 doi:10.1016/j.bbamcr.2021.118954

Sheng Y, Ali RA, Heyland A. 2022. Comparative transcriptomics analysis pipeline for the meta-analysis of phylogenetically divergent datasets (CoRMAP). BMC Bioinformatics 23:415 doi:10.1186/s12859-022-04972-9

Silhacek D, Murphy C. 2008. Moisture content in a wheat germ diet and its effect on the growth of *Plodia interpunctella* (Hübner). J. Stored Prod. Res. 44(1):36–40 doi:10.1016/j.jspr.2006.03.004

Sipilȁ S, Narici M, Kjaer M, Pőllȁnen E, Atkinson RA, Hansen M, Kovanen V. 2013. Sex hormones and skeletal muscle weakness. Biogerontology 14:231–245 doi:10.007/s10522-013-9425-8

Soneson C, Love MI, Robinson MD. 2015. Differential analyses for RNA-seq: transcript-level estimates improve gene-level inferences. F1000Res. 4:1521 dio: 10.12688/f1000research.7563.1

Staerk J, Conde DA, Tidière M, Lemaître JF, Liker A, Vági B, Pavard S, Giraudeau M, Smeele SQ, Vincze O, Ronget V, Silva RD, Pereboom Z, Bertelsen MF, Gaillard JM, Székely T, Colchero F. 2025. Sexual selection drives sex differences in adult life expectancy across mammals and birds. Sci. Adv. 11(40):eady8533 doi:10.1126/sciadv.ady8433

Stamboulian M, Guerrero RF, Hahn MW, Radivojac P. 2020. The ortholog conjecture revisited: The value of orthologs and paralogs in functional prediction. Bioinformatics 36:i219–i226 doi:10.1093/bioinformatics/btaa468

Studer RA, Robinson-Rechavi M. 2009. How confident can we be that orthologs are similar, but paralogs differ? Trends Genet. 25(5):210–216 doi:10.1016/j.tig.2009.03.004

Tieland M, Trouwborst I, Clark BC. 2018. Skeletal muscle performance and aging. JCSM 9:3–19 doi:10.1002/jcsm.12238

Tower J. 2010. Heat shock proteins and Drosophila aging. Exp Gerontol 46(5):355–362 doi:10.1016/j.exger.2010.09.002

Trombetti A, Reid KF, Hars M, Herrmann FR, Pasha E, Phillips EM, Fielding RA. 2015. Age-associated declines in muscle mass, strength, power, and physical performance: impact on fear of falling and quality of life. Osteoporos Int 27(2):463–471

Ubaida-Mohien C, Lyashkov A, Gonzalez-Freire M, Tharakan R, Shardell M, Moaddel R, Semba RD, Chia CW, Gorospe M, Sen R, Ferrucci L. 2019. Discovery proteomics in aging human skeletal muscle finds change in spliceosome, immunity, proteostasis and mitochondria. eLife 8:e49874 doi:10.7554/eLife.49874

Vasilaki A, Jackson MJ, McArdle A. 2002. Attenuated HSP70 response in skeletal muscle of aged rats following contractile activity. Muscle Nerve 25(6):902–905 doi:10.1002/mus.10094

Vaughan L, Marley R, Miellet S, Hartley PS. 2018. The impact of *SPARC* on age-related cardiac dysfunction and fibrosis in *Drosophila*. Exp. Gerontol. 109:59–66 doi:10.1016/j.exger.2017.10.011

Velders M, Diel P. 2013. How sex hormones promote skeletal muscle regeneration. Sports Med 43:1089–1100 doi:10.1007/s40279-013-0081-6

Viña J, Sastre J, Pallardó F, Borrás C. 2003. Mitochondrial theory of aging: importance to explain why females live longer than males. Antioxid Redox Signal 5(5):549–56 doi:10.1089/152308603770310194

Wakefield CB, Coulson PG, Loudon L, Newman SJ. 2020. Latitudinal and sex-specific differences in growth and an exceptional longevity for the Maori snapper *Lutjanus rivulatus* from north-western Australia. Fish. Res. 230:105634 doi:10.1016/j.fishres.2020.105634

Wang X, Jiang Q, Song Y, He Z, Zhang H, Song M, Zhang X, Dai Y, Karalay O, Dieterich C, Antebi A, Wu L, Han JDJ, Shen Y. 2022. Aging induces tissue-specific transcriptomic changes in *Caenorhabditis elegans*. EMBO J 41(8):e109633 doi:10.15252/embj.2021109633

Wickham H. 2016. ggplot2: elegant graphics for data analysis. Springer-VerlagNew York.

Wu T, Hu E, Xu S, Chen M, Guo P, Dai Z, Feng T, Zhou L, Tang W, Zhan L, Fu X, Liu S, Bo X, Yu G. 2021. clusterProfiler 4.0: a universal enrichment tool for interpreting omics data. Innovation 2(3):100141

Wyckelsma VL, McKenna MJ. 2016. Effects of age on Na+,K+-ATPase expression in human and rodent skeletal muscle. Front. Physiol. 7:316 doi:10.3389/fphys.2016.00316

[dataset]* Xu A, Teefy BB, Lu RJ, Nozownik S, Tyers AM, Valenzano DR, Benayoun BA. 2023. Transcriptomes of aging brain, heart, muscle, and spleen from female and male African turquoise killifish. Sci. Data 10:695 dio:10.1038/s41597-023-02609-x

Yu G, Wang LG, Han Y, He QY. 2012. clusterProfiler: an R package for comparing biological themes among gene clusters. OMICS: J. Integr. Biol. 16(5):284–287

[dataset]* Yu Y, Zhao C, Su Z, Wang C, Fuscoe JC, Tong W, Shi L. 2014. Comprehensive RNA-seq transcriptomic profiling across 11 organs, 4 ages, and 2 sexes of Fischer 344 rats. Sci. Data 1:140013 dio:10.1038/sdata.2014.13

